# Transposable Element (TE) insertion predictions from RNAseq inputs and TE impact on RNA splicing and gene expression in *Drosophila* brain transcriptomes

**DOI:** 10.1101/2024.06.07.597839

**Authors:** Md Fakhrul Azad, Tong Tong, Nelson C. Lau

## Abstract

Recent studies have suggested that Transposable Elements (TEs) residing in introns frequently splice into and alter primary gene coding transcripts. To re-examine the exonization of TEs into protein-coding gene transcripts, we re-analyzed a *Drosophila* neuron circadian rhythm RNAseq dataset and a deep long RNA fly midbrain RNAseq dataset using our Transposon Insertion and Depletion Analyzer (TIDAL) program. Our TIDAL results were able to predict several TE insertions from RNAseq data that were consistent with previous published studies. However, we also uncovered many discrepancies in TE-exonization calls, such as reads that mainly support intron retention of the TE and little support for chimeric mRNA spliced to the TE. We then deployed rigorous gDNA-PCR and RT-PCR procedures on TE-mRNA fusion candidates to see how many of bioinformatics predictions could be validated. By testing a *w1118* strain from which the deeper long RNAseq data was derived from and comparing to an *OreR* strain, only 9 of 23 TIDAL candidates (<40%) could be validated as a novel TE insertion by gDNA-PCR, indicating that deeper study is needed on using RNAseq as inputs into current TE-insertion prediction programs. Of these validated calls, the RT-PCR results only supported TE-intron retention. Lastly, in the *Dscam2* and *Bx* genes of the *w1118* strain that contained intronic TEs, gene expression was 2-3 times higher than the *OreR* genes lacking the TEs. This study’s validation approach indicates that chimeric TE-mRNAs are infrequent and cautions that more optimization is required in bioinformatics programs to call TE insertions using RNAseq datasets.

## INTRODUCTION

Transposable Elements (TEs) are insertional mutagens making up major fractions of animal genomes, yet we are still determining how TE mobilization affects gene expression, chromatin accessibility, and physiology. In multicellular organisms, most genes are filled with introns that are spliced out, thus introns are generally safe harbors for transposons to insert into without overtly disrupting the protein-coding exons. Since the spliceosome can effectively and precisely splice out introns varying widely in size, the impact of TEs on a gene’s regulation is hard to define if the TEs’ sequences are invisible to the splicing machinery. If an intronic TE’s sequences mutate so that alternative splicing can fuse the original gene exons to TE sequences, this ‘exonization’ event could be a genetic mechanism to incorporate TEs into novel mRNA isoforms [1, 2]. However, the extent of TE exonization in organism is still unclear because the field lacks extensive molecular validation of genomics predictions of chimeric TE-mRNA fusion transcripts arising from the revolution in high-throughput DNA and RNA sequencing.

To approach this question, we first consider a null hypothesis that perhaps most TEs residing in introns are not being exonized but rather acting as passive sequence platforms, which include evolving a selectable function for the binding of a transcription factor that connects the host gene to a new regulatory network [2–4]. A competing hypothesis is that TE exonization is frequent, and this proposition was recently examined by three studies [5–7] that devised custom bioinformatics programs to search transcriptome datasets from a lab stock and wild natural collections of *Drosophila melanogaster*.

The first study [7] developed a custom bioinformatics pipeline called TE-chim to search for TE-mRNA fusion reads from RNAseq data, and they posited that mRNA splicing to TEs to generate chimeric transcripts was a frequent event in the midbrain of the *Drosophila w1118* strain. A second study argues that ∼19% of somatic transcripts for TE-gene chimeric transcripts and could even make up 43% of transcriptomes in *Drosophila* [5]. A third study focusing on RNAseq datasets from *Drosophila* ovaries had a more conservative measure of ∼1% of transcriptomes displaying such TE-gene chimeras [6].

In all three studies claiming TEs are frequently expressed as parts of chimeric mRNAs with genes, there was minimal experimental support beyond the RNA sequencing datasets. Some RT-PCR amplicons were displayed in Oliveira et al [6] without amplicon sequencing confirmation or gDNA-PCR validation. The Coronado-Zamora and Gonzalez study [5] was only based on genomics bioinformatics and lacked PCR validation. Because these two studies primarily utilized wild *Drosophila* collections to highlight the diversity of TE landscapes and differential contribution of TEs to nearby genes, the ability to distribute and reproduce these natural *Drosophila* isolates is restricted. The Treiber and Waddel study [7] did not have any additional gDNA-PCR and RT-PCR experiments to back up the bioinformatics predictions from the RNAseq data; however the dataset came from a *w1118* strain derivative. Since *w1118* is one of the most widely used *Drosophila* lab strains, we followed this tractable opportunity to apply rigorous experiments to validate TE-gene chimeric events predicted by bioinformatics programs.

Our understanding of how intronic TEs impact a gene’s transcript maturation remains incomplete because standard DNAseq and RNAseq analysis algorithms are optimized to map sequencing reads to a reference genome and transcriptome, whereas intronic TEs that are either novel or even in a reference will not usually be included in gene models. Therefore, we developed our own bioinformatics algorithms for analyzing deep sequencing data for novel TE insertions and we have shown that TE landscapes are extremely diverse amongst *Drosophila* strains [8]. Our Transposon Insertion and Depletion Analyzer (TIDAL) program was originally designed around Whole Genome Sequencing (WGS) DNAseq datasets as inputs and is tuned to maximize specificity over sensitivity [8].

TIDAL’s stringent filters uses these cutoffs: (1) requiring split reads on both insertion junctions of a novel TE insertion; (2) prioritizing the symmetry of the split reads around the reference genome insertion site; (3) discarding split reads that still contained repetitive signatures from a low BLAT score; and (4) enabling the program to return all the split reads from a TE insertion call so that a user can apply an orthogonal query of the split reads on the *Drosophila* genome from the UCSC Genome Browser. We validated TIDAL’s specificity by testing 49 predicted TE-insertions that we could validate up to 88% of these events with gDNA-PCR [8, 9]. Although TIDAL does not put out as many candidate TE insertions predictions as other tools that promote higher sensitivity (i.e. [10–14]), we have greater confidence that TE insertions called in WGS by TIDAL can be confirmed by gDNA-PCR.

However, the next frontier would be to examine how TIDAL can handle RNAseq data as an input to primarily search for TE insertions in transcribe regions of the genome. Since RNAseq datasets are much more numerous and diverse from model organisms to human clinical samples, there could be good potential to leverage bioinformatics analysis of RNAseq data to discover novel TE landscape. Additionally, we hypothesized that TIDAL could also be a respectable benchmarking tool to examine the recent studies arguing that TE-gene exonization events could be a frequent event in *Drosophila* transcriptomes. The compact genome size and superb gene and TE annotations of *Drosophila melanogaster*, and the easy availability of lab strains makes these benchmarking studies on this model organism particularly attractive.

Thus, in this study we applied TIDAL to two *Drosophila* head and brain RNAseq datasets [7, 15] that have exceptionally deep RNA sequencing coverage that would not present a limitation compared to WGS. Furthermore, using the *Drosophila* lab strains like *w1118*, we could empirically test the confidence of TE insertion calls from RNAseq data with gDNA-PCR and RT-PCR to assess how frequently TE-gene chimeras accumulate compared to the standard gene transcripts. Our careful approach to TE-insertion validation shows that TE-gene chimeric transcripts are still very low in frequency amongst *Drosophila* brain/head transcriptomes, and while there is good promise to utilize TIDAL to find novel TE insertions from RNAseq data, our program and perhaps other programs as well still need to contend with unappreciated artifacts from RNAseq data that can reduce the confidence in valid TE insertion calls.

## RESULTS

### Considering RNAseq data as input for TIDAL TE insertion predictions

If intronic TEs were commonly exonized, we would envision two scenarios for how the RNAseq read coverage would diverge from canonical intron splicing (**Figure 1A**). If the TE sequence is seamlessly spliced to a gene’s exons, there should be RNAseq reads that span a seamless junction between the exon to the TE sequence. Alternatively, there could be transcription from the host gene promoter or autonomous transcription initiation from promoters within the intronic TE, and this scenario of intron retention would be reflected by RNAseq reads that span the intron-TE junctions.

**Figure 1.**
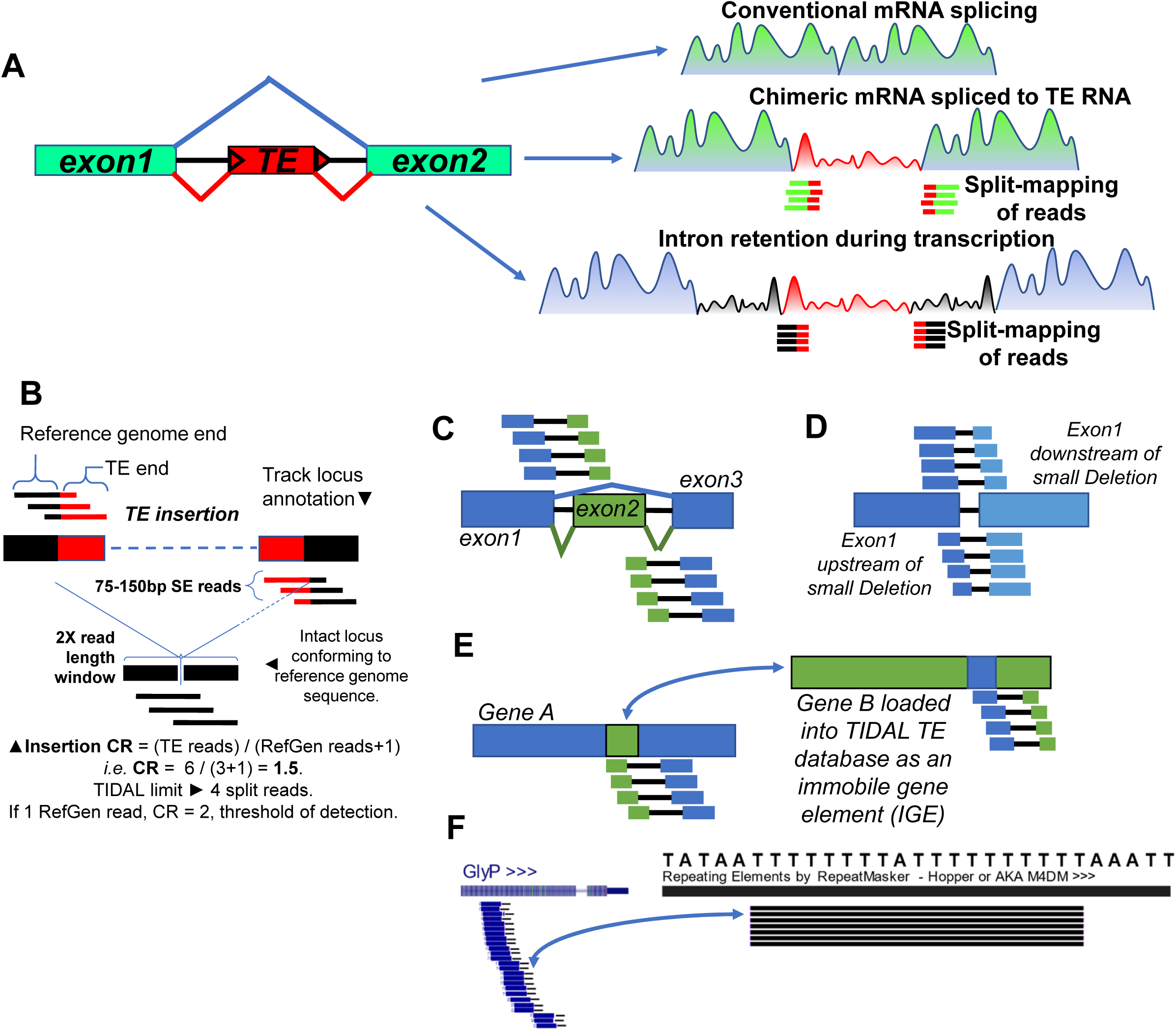
The chimeric TE-mRNA concept and TIDAL implementation using RNAseq data as input. (A) Diagrams considering how read coverage would reflect canonical exon splicing versus a TE-gene chimera versus intron retention during transcription of the intronic TE. Split reads representing these de novo TE insertions would not be mapped to a reference genome and transcriptome, requiring a specialized bioinformatics program like TIDAL and others. (B-E) Diagrams of TIDAL implemented on RNAseq to detect TEs in (B), Alternative splicing isoforms (C), small deletions like InDels (D), and potential gene-fusions that are more likely artifacts of similarity in sequences between different genes because some genes are loaded into TIDAL as an IGE (immobile gene element) (E). Depicted are split reads being aligned to the genomic structural variant by scripts within TIDAL. (F) A common artifact of a simple T-repeat sequence from reverse-transcribing from the Poly-A during Alternative Poly-Adenylation (APA) of a *Drosophila* gene like GlyP, where this simple T-repeat is part of the *hopper/M4DM* TE sequence. This additional simple-polynucleotide filter was added to TIDAL runs on RNAseq data.

We introduce these concepts because TIDAL and other *de novo* TE insertion prediction programs [8, 10–14] use split-mapping of RNAseq reads where one end must map to unique sequence in the reference genome and the other end must map to a database of TE consensus sequences. In the TIDAL algorithm design, split-mapping of DNAseq reads follows the regimen portrayed in Fig. 1B [8]. In a later version of TIDAL [16], we added a similar control feature of genes added into the repeats database to act as “Immobile Genetic Elements” (IGEs) as described in [7]. These IGEs help measure a false prediction rate in TIDAL that was below 12% in WGS DNAseq [16].

However, when we examined *Drosophila* head RNAseq data as input into TIDAL, the excessive number of IGEs being flagged in TIDAL outputs revealed a complexity in split-mapping of RNAseq reads that was not apparent in DNAseq reads. We will discuss the frequencies and fractional proportions of these IGE idiosyncrasies further below, but in Fig. 1C–F we first introduce these idiosyncrasies as diagrammed concepts. Because TIDAL splits a read to map each end within a window size (Fig. 1B), many IGEs can trick a TIDAL call if the reads span across the splicing of tiny introns (i.e. <100nt, Fig. 1C, **Supplemental Figure S1**) and genomic structural variants (SVs) like small insertions and deletions (InDels, Fig. 1D, S1A).

If a WGS library is properly prepared with complete shearing and sampling of the entire genome, the sequence diversity is immense and thoroughly distributed across the multitude of reads. Transcriptomes, however, may only represent <10% of the entire sequences of an animal’s genome, with different expression levels that can bias many gene sequences over others, and different protein-coding genes can share short similar sequences if they are encoding a commonly shared protein domain. To fully consider RNA maturation steps, we conjecture that short sequence compositions within RNAseq inputs are influencing the significant differences in TIDAL outputs compared to WGS DNAseq inputs.

Furthermore, the input of RNAseq into TIDAL also raised the calls of two distinct genes that might appear to form a two-mRNA fusion transcript (Fig. 1E) whereas these types of calls were low with DNAseq inputs. The TIDAL program [8] was originally designed to take as input WGS DNAseq from Illumina reads as short as 50 nucleotides (nt), and uses the Bowtie v1 algorithm [17] for split-mapping of each read’s end using just 22nt but allowing up to 3 mismatches. Given how transcriptomes sequences are just more naturally biased in short sequences that could be similar between genes compared to entire genomes (Fig. S1B), the frequency of finding these kind of shared alignments between disparate genes became quite high. We favored this interpretation over the other molecular possibility of artifactual mis-priming of distinct gene amplicons during PCR amplification steps of RNAseq library construction [18, 19].

Lastly, this short read-alignment mismatch problem resulted in TIDAL detecting an abnormally high number of putative insertions of the *hopper*/*M4DM* TE in genes (Fig. 1F). In the two-gene fusion issue illustrated in Fig. S1B, we noticed a pattern of a simple sequence like Poly-T in the *Zelda* gene, this could have been generated from reverse-transcription of a Poly-A tail during library preparation. There is also a Poly-T sequence within the *hopper/M4DM* TE sequence that TIDAL was latching onto to calling an excessive number of false positives. WGS DNAseq libraries are immune from this Poly-T artifact that is easy to see being formed in RNAseq libraries. We tweaked TIDAL for RNAseq to ignore this Poly-T artifact to cut down on false *hopper/M4DM* calls, but the other simple shared sequences between two distinct genes were too numerous and diverse for us to develop a suitable filter screen away these two-gene artifacts, although the annotation information within TIDAL output tables allows us to mark and count these two-gene artifacts.

### TE-mRNA fusion calls are a minority of TIDAL prediction events from Drosophila neurons and brain RNAseq inputs

We first tested TIDAL’s functionality on RNAseq by inputting high-quality transcriptomes from four purified sets of *Drosophila* neurons repeatedly sampled across 6 circadian time points [15]. These 48 RNAseq libraries enabled us to examine how reproducible or variable were potential TE-mRNA fusion calls by TIDAL since these neurons should have at least a large backdrop of consistently expressed genes with a subset of circadian oscillating genes. We also tested a very-deep longer-read (250x250PE) RNAseq dataset from the Treiber and Waddell study that proposed frequent TE exonization events and non-autonomous TE transcript expression in *Drosophila* midbrains [7].

We tracked the variation in library sequencing depths and read-mapping proportions to the *Drosophila* reference genome for each of the 48 circadian rhythm *Drosophila* neurons RNAseq libraries (**Figure S2A**). We also merged each of the 12 libraries into their single neuron type (LNd, LNv, DN1, and TH, Fig. S2B), and after analyzing all these RNAseq datasets as inputs into TIDAL, we then counted the TIDAL outputs by three categories: (1) TE-mRNA fusions as our main feature of interest, (2) a genuine molecular event of InDels and Splicing isoforms that activate a TIDAL call, and (3) artifacts of two mRNAs being called as a fusion event (**Figure 2A**, Fig. S2A). Regardless of variations in library reads depth and genome-mappable read fractions in these circadian rhythm RNAseq datasets, two-mRNA fusions were the major proportion of TIDAL calls followed by InDel/Splicing isoform calls. TE-mRNA fusion calls were the minority, between 18-22% in these TIDAL calls.

**Figure 2.**
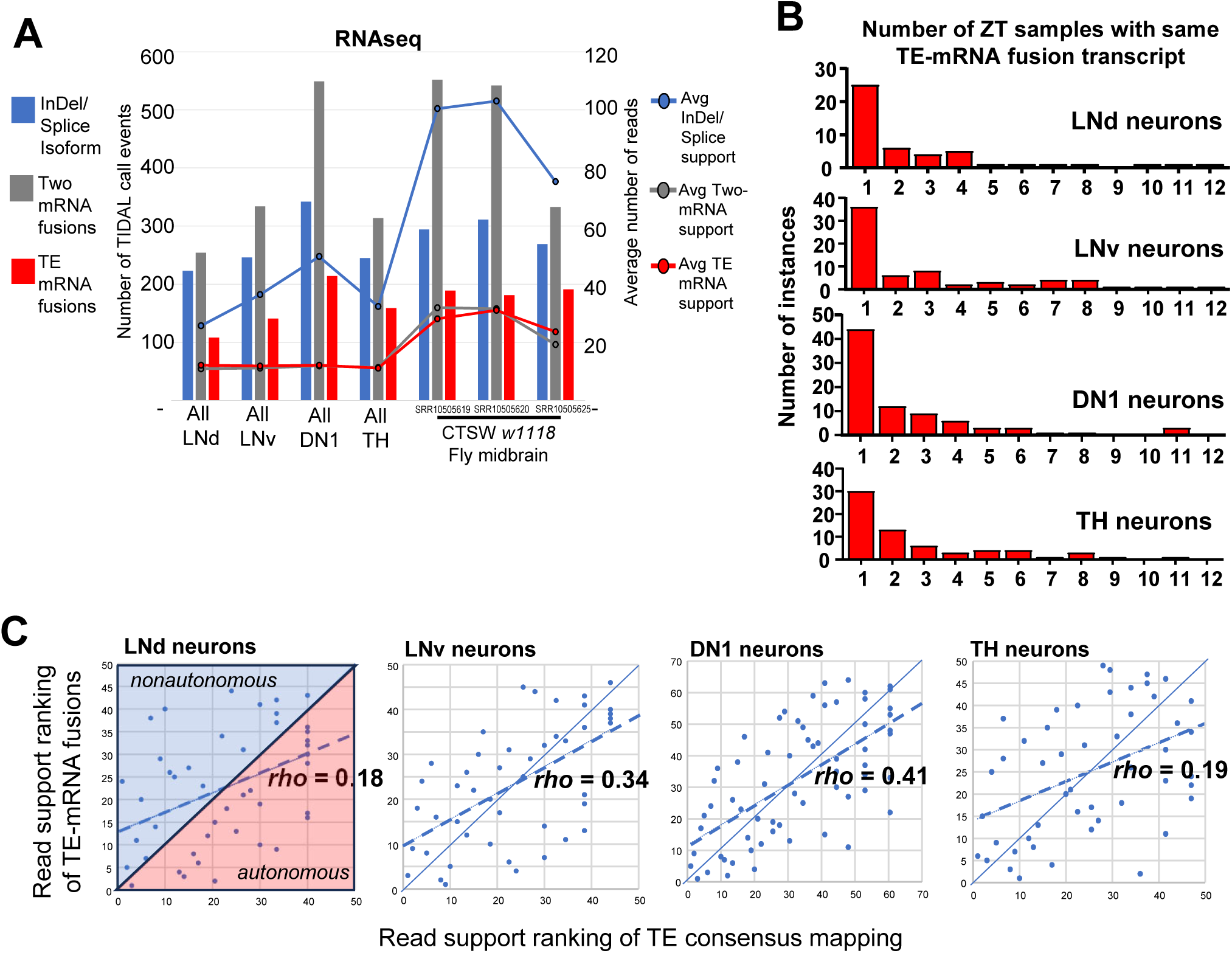
Comparing TE-mRNA fusions and insertion calls versus other TIDAL calls from *Drosophila* brain RNAseq inputs. (A) Histograms of the number of the Zeitgeiber Time (ZT) samples supporting the same consistent TE-mRNA fusion calls by TIDAL in the circadian rhythm RNAseq dataset. (B) How RNAseq inputs perform in TIDAL with the bar graph shows the number of TIDAL calls (left Y-axis number) for events representing an InDel/Splice isoform, a two-mRNA fusion call, or a TE-mRNA fusion call. Lines in the bar graph display the average number of reads (Right Y-axis) that support the different TIDAL calls. (C) Scatterplots show some positive correlation between read support ranks of TEs in TE-mRNA fusions versus consensus sequence mapping events. The diagonal-splitting lines of each scatterplot separates the non-autonomous TEs expression tied to the gene in the upper-diagonal half versus the autonomous TE expression dots that are in the lower-diagonal half.

These circadian neuron RNAseq libraries have 12 timepoints that can serve as replicates to ask how many times a given TE-mRNA fusion call is seen repeatedly, which would lend support that the call is a true positive. The frequency histograms show that the vast majority of the TE-mRNA fusion calls are only seen in a single timepoint sample in this circadian RNAseq dataset (Fig. 2B). An independent RNAseq dataset from the *Drosophila* midbrain that was run through TIDAL also exhibited just 18-24% of the calls representing TE-mRNA fusion events, while two-mRNA artifacts also dominated in these TIDAL calls. Although TIDAL outputs enable simple detection and filtering of the false-positive Two-mRNA fusion events, we realize that TE-mRNA fusion events need to be inspected further.

To re-examine whether the recent claims of TE expression being non-autonomous from host genes in *Drosophila* brain RNAseq [7] would be supported by our analysis, we performed a correlation analysis between the rankings by the counts of read-support for TEs in the consensus element counts versus TE-mRNA fusion calls (Fig. 2C), which is a similar but different analysis as the Figure 6A in the Treiber and Waddell study [7]. With an overall consistent positive correlation of TE-mRNA fusion call read support and TE-consensus coverage read support, there were several TEs in our analysis that were in the upper-diagonal half of these scatterplots, consistent with non-autonomous TE expression as depicted in the scatterplot in Fig. 6A in the Treiber and Waddell study [7]. However, in contrast to the claim that all TE expression was non-autonomous, our analysis showed many TEs were expressed autonomously from the mRNA (lower-diagonal half), and this would explain the relatively low (<42%) *rho* correlation coefficients. Our results indicate that TE-mRNA fusion events need more scrutiny to their validity than what the bioinformatics predictions may indicate.

### TIDAL performance differences between RNAseq and WGS-DNAseq as inputs

Since TIDAL only calls TE-mRNA fusion events as a minority of the outputted predictions from RNAseq inputs, we compared a set of recent WGS-DNAseq libraries analyzed under the same version of TIDAL [16]. Despite an average ∼50% fewer reads for each TIDAL event call compared to the RNAseq inputs, the WGS-DNAseq inputs yielded the overwhelming majority of events were TE insertions into gene-proximal loci (**Figure 3A**). As expected from the biochemistry of WGS-DNAseq library preparation, calls for InDels were miniscule. The major artifact of Two-mRNA fusions making up most of TIDAL calls from RNAseq inputs was many-fold lower in the WGS-DNAseq inputs. These results reaffirm the robustness of TIDAL’s TE-insertion call outputs when WGS-DNAseq is used as inputs.

**Figure 3.**
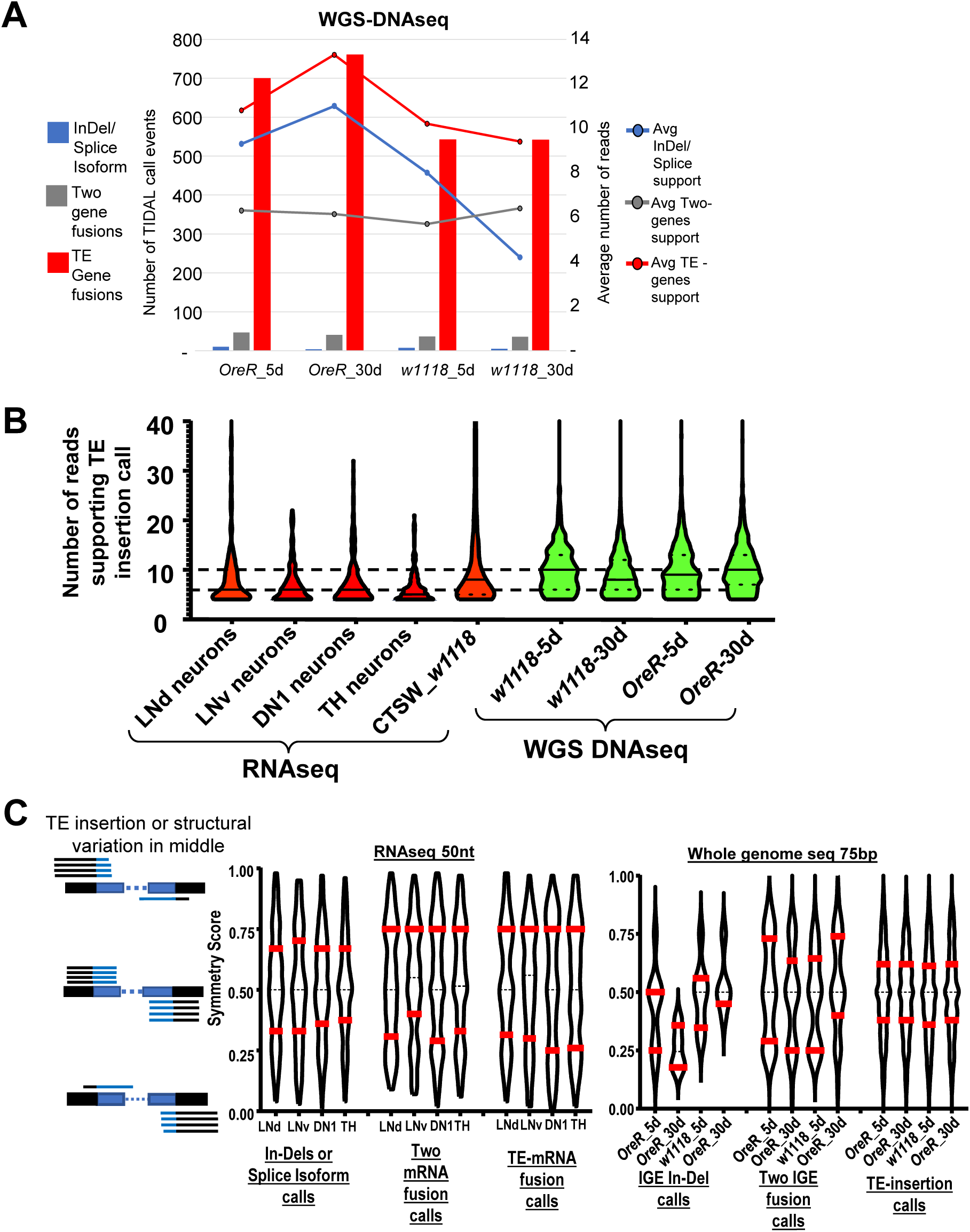
TIDAL performance differences between RNAseq and WGS-DNAseq as inputs. (A) Similar bar and lines graph as Fig. 2A but for WGS-DNAseq as the standard input into TIDAL. The TE-gene fusions in the WGS-DNAseq is a much greater proportion even though there is a lower number of supporting reads. (B) Violin plots showing a more skewed bias in RNAseq inputs (colored red) of generally fewer split reads supporting each event call, whereas DNAseq inputs (colored green) have a more balanced distribution of more split reads supporting each event call. TIDAL requires at least 4 reads that span the structural variation breakpoint to make a call, so all of these violin plots have a wide base at 4 reads. (C)Violin plots of the distribution of symmetry scores for TIDAL calls of genome structural variants from *Drosophila* neuron RNAseq versus whole fly WGS libraries. Red bars mark quartiles, the dashed midline is the mean.

TIDAL was originally designed on using WGS-DNAseq Illumina sequencing reads as inputs, and the specificity of choosing valid calls was tuned with several quality filters [8]. On quality filters are requirements such as (1) a minimum of 4 split reads of support covering the TE insertion junction, (2) that the ends of the split reads have a BLAT score [20] of >50% to catch some of the unmasked and unannotated simple-sequence repeats, and (3) that these reads are spread out within the defined window size of 2X times the read length (Fig. 1B). The distribution of these split reads on either side of the TE insertion also allows us to calculate a “symmetry score” for each TE insertion call, such as a 50% score that is ideal symmetry (half reads on each side of the insertion), and scores towards the extremes of 1% and 99% exhibit biased distribution of the insertion-spanning reads.

When we compared these quality filter scores of TE insertion calls between RNAseq and WGS DNAseq inputs, the violin plots showed that the majority of TE-mRNA fusion calls in RNAseq were skewed at the low 4-read minimum (Fig. 3B), with the mean at just 6 supporting reads. In contrast, the distribution of the number of supporting reads for WGS-DNAseq was more balanced with the mean of 10 supporting reads for TE-insertion calls. There was also an extended upward tail of varying number of reads supporting TE insertion calls, reflecting aspects of non-uniform read coverages in both RNAseq and WGS-DNAseq libraries that we currently do not yet fully understand.

To illuminate the performance differences between these two types of high-throughput sequencing inputs, we compared symmetry scores distributions for each of the event calls made by TIDAL for RNAseq versus WGS-DNAseq (Fig. 3C). We expect InDels/Splicing events in RNAseq to be a robust molecular feature for mature RNA transcripts, and the violin plots of the symmetry scores for InDels/Splicing events fit an archetypal shape – a central bulged mean at 50% and the quartiles nestled closer to this central mean. This archetypal violin plot shape was also shared in the TE-insertions calls’ symmetry scores in the WGS-DNAseq outputs, which have strong confidence in the validity of these event predictions. The presumptive artifacts such as the Two-mRNA fusions in both inputs displayed deviating violin plot shapes from archetypal shapes of InDels/Splicing events in RNAseq and TE-insertions in WGS-DNAseq, suggesting that this parameter and others may be useful to better screen out artifacts.

### PCR validation of predicted TE insertion in Drosophila genomes from RNAseq inputs

To test whether mRNA-TE chimeras are frequently present in *Drosophila* brain transcriptomes as proposed in previous studies [5–7], we developed rigorous gDNA-PCR and RT-PCR approaches to assay gDNA and total RNA from *Drosophila w1118* and *Oregon-R* (*OreR*) heads. We selected 19 predicted TE-gene insertions that both TIDAL and TEchim predicted from the same *w1118* mid-brain RNAseq inputs (**Table 1**). We also selected 3 cases and 4 cases only predicted by TEchim or only by TIDAL, respectively. The summary of our gDNA-PCR results is tabulated in Table 1 and will be elaborated upon in the text discussion below.

**TABLE 1.**
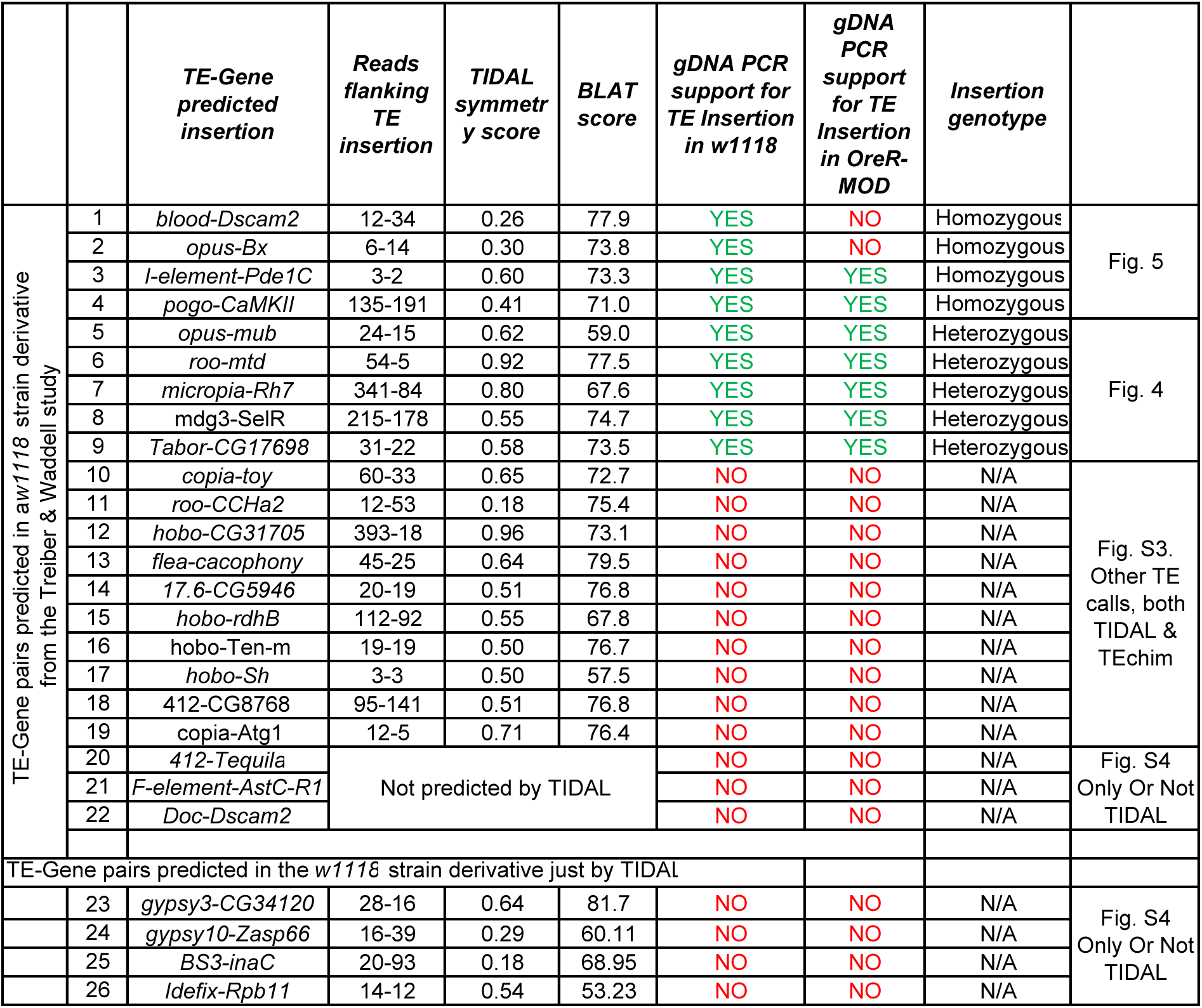
List of TE-Gene pairs evaluated by PCR in this study.

For each of these 26 cases, we conducted gDNA-PCR analysis with a series of primers to first validate if the predicted TE was indeed inserted into the gene regions in *w1118* and *OreR* strains genomic DNA. These panels consist of an amplicon called GPCR1 that are made by gene-specific primers the immediately flank the putative TE insertion, and amplicons GPCR2 and GPCR3 that use one primer that is gene-specific and the other primer binds to the adjacent terminus of the putative TE insertion (**Table S1, Figure 4**). When a TE insertion is true and too large for the GPCR1 primers to yield an amplicon (*i.e.* a full-length *opus* insertion is 7.5kb [21], so GPCR1 primers 2-5 can only amplify the allele lacking the TE, Fig. 4A-iii), then the GPCR2 (primers 1-3) and GPCR3 (primers 4-5) amplicons are short enough to be efficiently amplified to confirm the TE inserted into the gene intron.

**Figure 4.**
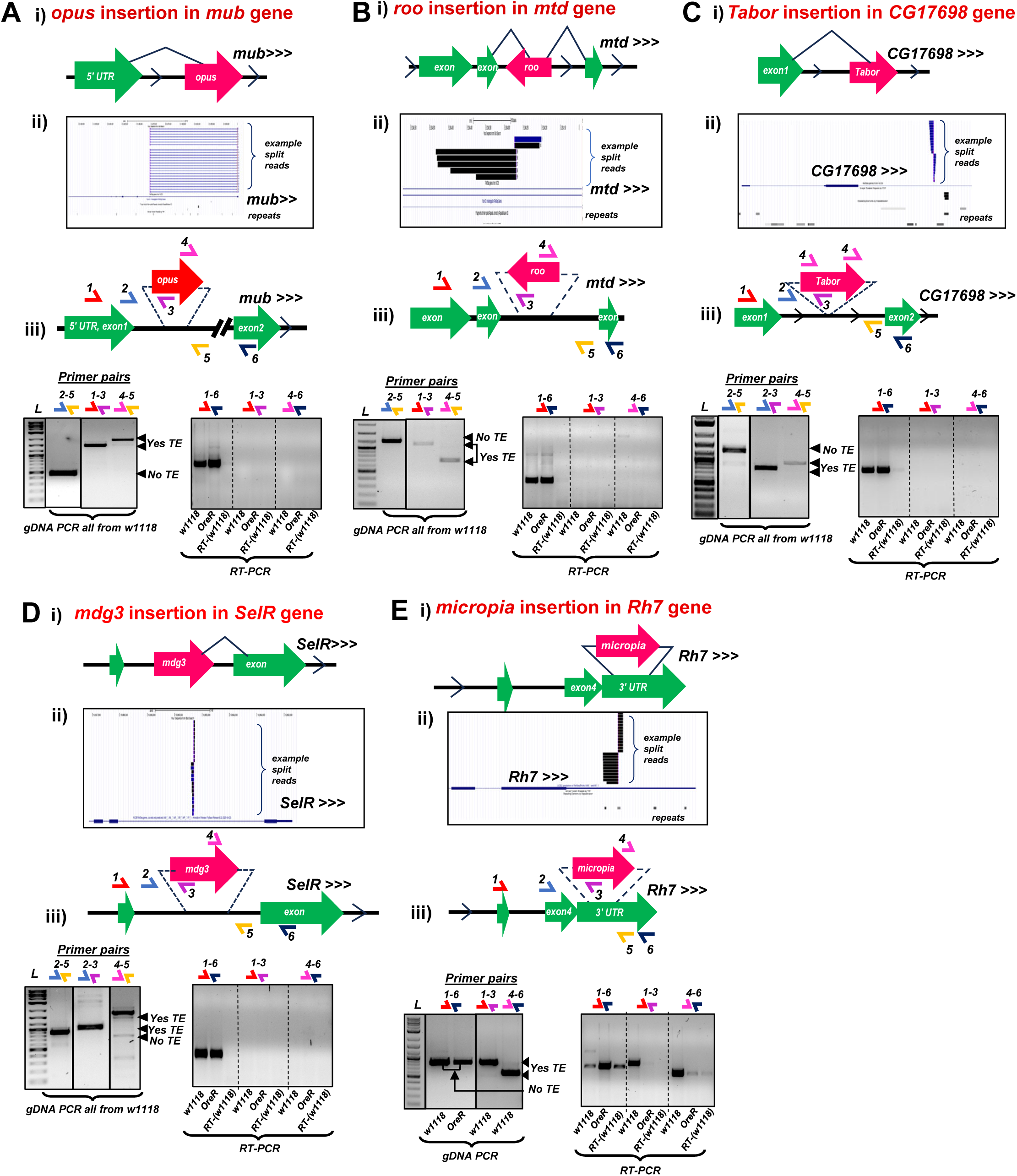
Rigorous gDNA-PCR validation of heterozygous TE insertions into genes in the *Drosophila w1118* strain genome. Panels correspond to the TE-gene pairs (A) *opus*-*mub,* (B) *roo*-*mtd,* (C) *Tabor-CG17698,* (D) *mdg3-SelR,* and (E) *micropia-Rh7,.* Each figure panel is divided in parts (i) that is a diagram of the presumptive TE-gene splicing event proposed by Treiber and Waddell 2020, (ii) UCSC Genome Browser snapshots of the example split reads support for the TE insertion from TIDAL analysis of the *w1118* midbrain RNAseq data, (iii) gel images of gDNA-PCR amplicons (left set) and RT-PCR amplicons (right set) from the various sets of primer pairs illustrated in the diagram above the gel images. Solid lines around gels marked cropped gel images; dashed lines are demarcating lane sections on a single gel.

This experimental approach allowed us to confirm five TE insertions predicted by TEchim and TIDAL from RNAseq inputs, and our data show they are heterozygous in the *w1118* strain genomes (Fig. 4A-E). These TEs are *opus* insertion in the intron of *mub* (Fig. 4A), *roo* in an intron of *mtd* (Fig.4B), *micropia* in an intron of *Rh7* (Fig.4C), *Tabor* in the intron of *CG17698* (Fig.4D), and *mdg3* inserted in a *SeIR* intron (Fig.4E). We conclude these TE insertions were heterozygous because the GPCR1 amplicon could only amplify the wild-type allele lacking the TE, whereas GPCR2 and GPCR3 amplicons were detected and confirmed for the TE insertion by amplicon sequencing. All five of these heterozygous TE insertions were also observed in the *OreR* strain (Table 1, **Figure S3**).

With four other TE insertions also predicted by TEchim and TIDAL from RNAseq inputs, our data showed these were homozygous TE insertions in the *w1118* strain genomes (**Figure 5A-D**). We confirmed a *pogo* inserted in the *CaMKII* gene intron (Fig.5A), *I-element* in the *Pde1C* gene intron (Fig.5B), *opus* inserted into the *Bx* gene intron (Fig.5C), and *blood* inserted into the *Dscam2* gene intron in *w1118* flies (Fig.5D). These TE insertions were homozygous in the *w1118* genome because a theoretical amplicon for GPCR1 primers was too long to be efficiently amplified, but GPCR2 and GPCR3 amplicons that were sequenced confirmed the TE insertion identity. Two of these homozygous TE insertions in *w1118* were also present and homozygous in *OreR* (Table 1, Fig. S3). We interpret that these seven TE insertions commonly shared between *w1118* and *OreR* strains were in the ancestral *D. melanogaster* strain, yet it is mysterious how five of these TEs can be maintained as heterozygous alleles when *Drosophila* transvection and meiotic recombination have not driven the allelic conversion to the homozygous TE insertion state [22, 23].

**Figure 5.**
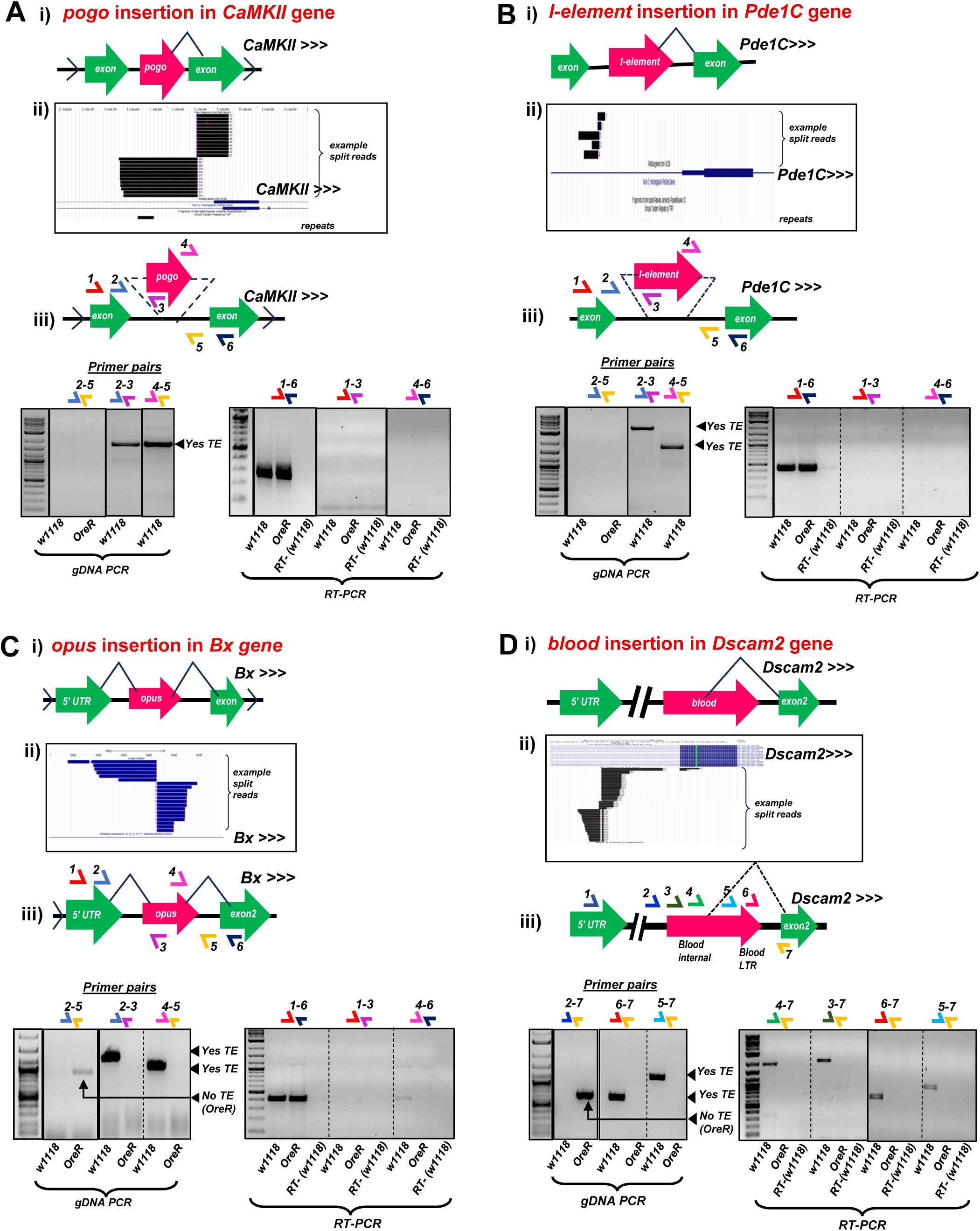
Rigorous gDNA-PCR validation of homozygous TE insertions into genes in the *Drosophila w1118* strain genome. Panels correspond to the TE-gene pairs. (A) *pogo*-*CaMKII,* (B) *I-element-Pde1C,* (C) *opus-Bx,* and (D) *blood-Dscam2.* Each figure panel is divided in parts (i) that is a diagram of the presumptive TE-gene splicing event proposed by Treiber and Waddell 2020, (ii) UCSC Genome Browser snapshots of the example split reads support for the TE insertion from TIDAL analysis of the *w1118* midbrain RNAseq data, (iii) gel images of gDNA-PCR amplicons (left set) and RT-PCR amplicons (right set) from the various sets of primer pairs illustrated in the diagram above the gel images. Solid lines around gels marked cropped gel images; dashed lines are demarcating lane sections on a single gel.

Of the 17 remaining TE insertion predictions, our gDNA-PCR experiments were unable to confirm an actual TE insertion in the intron of the host gene in neither *w1118* nor *OreR* (Table 1). Although four and three putative TE insertions were only predicted by either TIDAL or TEchim, respectively, the other nine of these TE insertion predictions were predicted by both programs, and we noted the relatively high number of supporting split reads that appeared to flank the TE insertion and displayed ideal TIDAL symmetry scores and BLAT scores. The negative results of testing for TE insertions predicted by both TEchim and TIDAL are shown in (**Figure S4**), while TE insertions only predicted by either TIDAL or TEchim are shown in (**Figure S5**). Only the GPCR1 amplicon indicated a wild-type genotype at each gene intron location, with no GPCR2 or GPCR3 amplicons detected. These results indicate the majority (∼65%) of the predicted TE insertions using RNAseq as inputs could potentially be false positives that will require deeper study because our current TIDAL metrics (high supporting split reads, symmetry scores, and BLAT scores) cannot yet distinguish between the 9 true-positive TE insertions from the 17 false-positive predictions.

### Effect of transposon on host gene RNA splicing and steady state mRNA accumulation

We utilized the similar primers that validated the 9 TE genomic insertions to examine for possible TE-mRNA chimera formation in an RT-PCR assay. In the rightmost subpanel of Fig. 4Aiii to 4Diii and Fig. 5Aiii to 5Ciii, the RT-PCR results only showed a robust amplicon corresponding to the complete canonical splicing of the two exons flanking the intron harboring the TE. Other primer pair amplicons designed to detect a potential TE-mRNA chimera repeatedly failed to detect a clear sign of the putative TE-mRNA chimera proposed by Treiber and Waddell [7]. Notably, all the RNAseq split reads that informed TIDAL to also make the same TE insertion prediction as TEchim as displayed in Fig. 4 and Fig. 5 are only mapping to the intronic RNA directly flanking the TE insertion. The lack of any split reads in the TIDAL outputs between the TE splicing into the host gene’s exons is supported by our negative results of few clear amplicons corresponding to the TE-mRNA chimera.

Only two TE insertions in genes yielded amplicons supporting a TE-mRNA chimera. There is a true TE-mRNA fusion of *micropia* inserted in the 3’ UTR of the *Rh7* gene, but because this is an exon, splicing did not contribute to this TE-mRNA chimera. In addition, this TE does not affect the *Rh7* open reading frame but might impact the mRNA’s expression level (Fig.4E). The *blood* TE is inserted into the first intron of the *Dscam2* gene in the *w1118* strain (Fig.5D), but contrary to the proposition by Treiber and Waddell [7] that *blood* was splicing frequently into the second exon, our RT-PCR results only detected transcription indicative of intron retention, not splicing (Fig. 5Diii). This RT-PCR result is consistent with the TIDAL outputs showing the vast majority of TE split reads between *blood* and *Dscam2’s* first intron are only supporting intron retention. Between the gDNA-PCR and RT-PCR results, there is little experimental support for validating TE-mRNA chimeras that are predicted from the analysis of RNAseq inputs.

Nevertheless, the polymorphic TE intronic insertions in the *Dscam2* and *Bx* genes in only the *w1118* strain but not in the *OreR* strain presented an interesting opportunity to test how a TE insertion would impact the efficiency of host mRNA splicing or steady level of mRNA accumulation. We designed a series of exon-specific primer pairs spanning the intron containing the TE in the *w1118* strain as well as additional exons separated by introns downstream (**Figure 6**). We then quantitated amplicon amounts normalized relatively to the *Rp49* housekeeping gene, and repeatedly observed 2-to-4-fold greater *Bx* and *Dscam2* transcript levels, respectively, in the *w1118* head RNAs compared to *OreR* fly heads (Fig. 6Aii, 6Bii). The relative quantitation methodology showing greater host gene mRNA expression in *w1118* compared to *OreR* was confirmed by an independent absolute quantitation methodology with droplet-digital PCR (ddPCR, Fig. 6Aiii).

**Figure 6.**
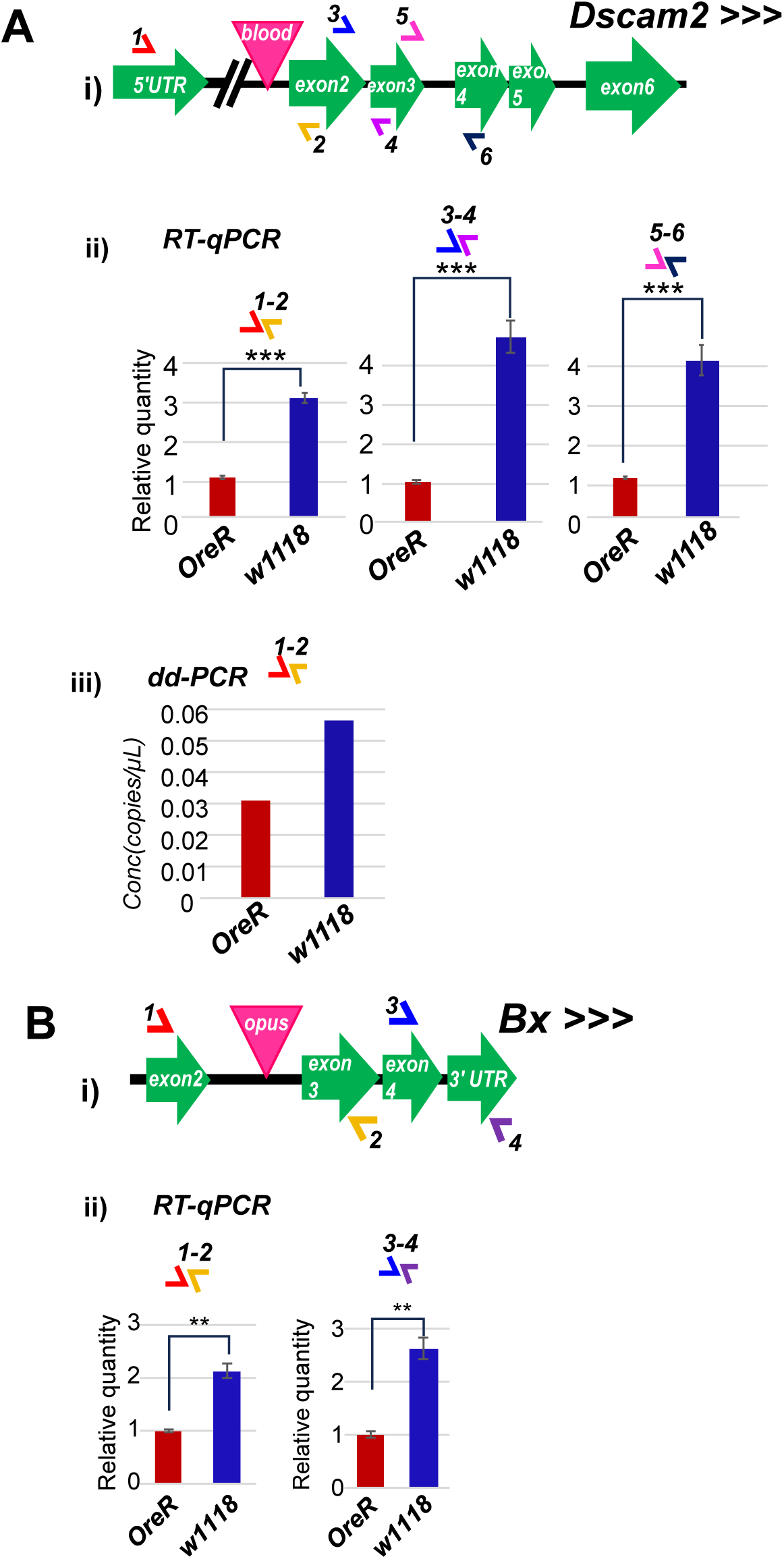
Testing the effect of a TE insertion within an intron on host gene mRNA splicing and expression. Diagrams showing the *w1118* strain has *blood* TE insertion in *Dscam2* gene (A-i) and *opus TE* insertion in *Bx* gene (B-i), while the *OreR* strain does not have those TE insertions in the corresponding genes. (A-ii) RT-qPCR assay examining spliced *Dscam2* mRNA levels in the heads of *w1118* and *OreR*. (A-iii) ddPCR assay replicating the RT-qPCR result in (A-ii). (B-ii) RT-qPCR assay examining spliced *Bx* mRNA levels in the heads of *w1118* and *OreR*. T-tests p-values: *=*p*<0.05, **=*p*<0.005, ***=*p*<0.001.

If the *blood* and *opus* TEs inserted into *Dscam2* and *Bx* introns, respectively, are full length, they each would expand the intron sizes by ∼7.5kb. We could not disentangle if the TE insertion was counterintuitively enhancing intron splicing, or if the TE was serving as an enhancer to increase transcription activation because downstream exon amplicons were just as elevated in *w1118* versus *OreR*. Our results clearly show absence of supporting evidence for alternative splicing that would generate TE-mRNA chimeras, but these two examples could represent the first cases of *Drosophila* TEs providing a regulatory element to enhance gene expression, similar to examples in mammals reviewed in [24, 25].

## DISCUSSION

Although recent studies [5–7] have claimed that TEs in *Drosophila* may be frequently splicing into host gene exons to generate TE-mRNA chimeras, the interpretations have mainly relied upon bioinformatics predictions without introspection of the supporting read patterns. Only one of the studies [6] performed some RT-PCR experiments on potential TE-mRNA chimeras, but there was no confirmation first by gDNA-PCR nor utilization of multiple primer pairs to ensure experimental rigor. The extensive gDNA-PCR and RT-PCR analyses in our study demonstrate the lack of evidence supporting TE-mRNA chimeric splicing. Thus, the question remains unanswered: how extensively do TEs affect host gene expression, via splicing or perhaps other mechanisms like transcription activation via acting like an enhancer platform [26]?

A transcriptome is a constrained subset of the genome’s entire sequence diversity, and this feature may be reflected in our study whereby RNAseq as inputs into bioinformatics programs like TEchim and TIDAL can lead to the same artifact predictions of TE insertions that cannot be validated by gDNA-PCR (Table 1). Currently, TIDAL faithfully detects TE insertions events using WGS-DNAseq data at a validation rate of >66% [8], whereas using RNAseq inputs lowers the validation rates to ∼40% (9 of 23 tested TE insertion calls). Our next efforts to update TIDAL will be to search for indicative patterns that will help us screen out artifactual two-mRNA fusion events to improve the confidence in TE insertion calls and TE-mRNA chimeras.

Our study challenges the notion that TEs splicing into mRNAs may not be widespread [5–7], and there is the unaddressed concern of imprecision with TE exonization which would generate many deleterious and non-functional transcripts turned over by Nonsense Mediated Decay (NMD) processes [27]. From a few studies in plants [28, 29], to one notable study in human cells with the ORF0 sequence in the TE LINE L1 5’ UTR [30], there can be instances where TE-mRNA chimeras could form, but these studies acknowledge NMD likely safeguards most deleterious transcripts from impacting host organism fitness.

In addition, the RNA interference (RNAi) pathway also silences TE expression in *Drosophila*, and we demonstrated that augmenting RNAi improves longevity by mitigating negative effects of increased TE expression in aged flies [16], a phenotype that has also been frequently observed in mammals [31–34]. We favor the idea that RNAi is mainly silencing TE transcripts that are autonomously expressed (Fig. 2C) without also negatively silencing many neighboring genes because TE-mRNA chimeras are infrequent.

However, we acknowledge that the literature includes several examples of TEs inserting into host genes and affecting gene expression positively to yield potentially novel functions. A *Doc* TE insertion in the coding sequence of *CHKov1* truncates CHKov1 mRNA to encode a shorter peptide that confers greater sigma virus resistance to *Drosophila* [35]. In another case, successive insertions of the *Accord, HMS-Beagle* and *P*-element TEs inserted at the *Drosophila* cytochrome P450 (*Cyp6g1*) gene improved resistance to the insecticide DDT [36]. Third, there is the KRABINER fusion gene of a mariner TE and KRAB domain protein modulating gene expression in bat cell cultures [37], but its true biological function in the bat animal remains unclear.

Lastly, TEs may exert regulation in *cis* to host genes via serving as a novel enhancer platforms that recruit new transcription factor binding situations [3, 4, 24, 26]. We are speculating this hypothesis could explain the influence the intronic insertion of *blood* and *opus* TEs into *Dscam2* and *Bx* neuronal genes, respectively, in just the *w1118* fly strains (Fig. 6). The functional consequence of these higher *Dscam2* and *Bx* mRNA levels in *w1118* flies compared to *OreR* is not yet clear, but perhaps these TEs provide new regulatory elements that enhances *Dscam2* and *Bx* mRNA expression in *w1118* flies compared to the control *OreR* flies that completely lack these TEs in its *Dscam2* and *Bx* genes. Future genetics and molecular biology experiments will improve our genomics and bioinformatics analyses of TEs’ impact on gene expression to uncover additional TE-gene interactions like the ones examined in this study.

## MATERIALS AND METHODS

### Accessing RNA-seq data sets

For this analysis we downloaded a publicly available RNA-seq dataset of *Drosophila* circadian rhythm cycling neurons [15] from NCBI Accession #GSE77451. This time-series data set contains 48 RNA samples extracted from 4 types of neurons in fruit fly brains, including dorsal lateral neurons (LNds), ventral lateral neurons (LNvs), dorsal neurons group 1 (DN1s), and dopaminergic neurons (TH). We also downloaded the full RNAseq data from the w1118 *Drosophila* midbrain from the Treiber and Waddell study [7] under the NCBI Accession #PRJNA588978.

### Detecting TE insertions using TIDAL and TEchim

We first used TIDAL program [8] to detect TE insertions for each sample and pooled samples for each type of neuron in Abruzzi et al [15] and in the *w1118 Drosophila* midbrain from the Treiber and Waddell study [7]. Specifically, TIDAL removed adaptors sequence, Poly-A, and low-quality bases from raw reads and duplicates. Then, pre-processed reads were aligned to the *Drosophila melanogaster* reference genome Release 6 (Dm6) and unmapped reads, which potentially contain de novo inserted TE segments, were kept. Viral RNA, structural RNA, Repbase sequence, and TE sequence were further removed.

Next, to identify the TE-gene junction, 22nt-long sequence at both 5’ and 3’ ends were taken from each read and mapped to TE consensus sequence and immobile gene elements (IGE) database and repeat-masked Dm6 reference genome. If one end was mapped to TE or IGE and the other end was uniquely mapped to reference genome, reads with such split short reads were kept. Then, reads within 300nt and with one end mapped to the same TE were clustered together and clusters with size larger than 4 reads were kept. To further reduce false positives, clusters with BLAT score >83% and span size smaller than (read length / 2) – 22nt were filtered. A metric called Coverage Ratio, which is the ratio between number of split reads containing TE and the number of Dm6 mapped reads plus a pseudo count, was also calculated for additional filtering [8]. We also implemented a screen to remove sequences against the *hopper* TE sequence.

TEchim [7] used a similar strategy to detect TE insertions. After pre-processing steps including splitting reads at both ends, TEchim mapped these in-silico paired-end reads to a repeat-masked reference genome combined with TE consensus sequence. Reads covering TE-gene breakpoints were kept as input for BLAST program for annotation. However, TEchim doesn’t have a step to remove adaptors like TIDAL has, so we used Trimmomatic software to remove adaptors from raw fastq files.

After running TIDAL and TEchim programs, we summarized unique TE insertions across all 12 samples for each type of neurons. Unlike TEchim, which only targets on finding TE insertion, TIDAL can also identify gene translocation and insertion/deletion variant. Therefore, we counted the number of unique insertions based on annotation of inserted sequence and its neighboring gene. We only utilized our TEchim run to confirm the results from the previously published Treiber and Waddell study [7], and to confirm the reproduced predictions listed in Table 1 and Table S1.

### Genomic DNA extraction and PCR

The *D. melanogaster w1118-iso* and *OreR-MOD* strains were used in this study. Both strains were raised at 25°C on standard cornmeal food. Genomic DNA were extracted from 7-day old 30 female fly heads using NEB Monarch Genomic DNA Purification Kit following the manufacturer provided protocol. DNA quantity and quality was checked on NanoDrop One Microvolume UV-Vis Spectrophotometer (ThermoFisher).

Genomic PCR was performed using NEB Phusion High-Fidelity DNA polymerase (GC buffer) with different primer combinations described in Table S2 with an input of approximately 50ng/ul DNA. Genomic PCR1 (GPCR1) was performed using a gene specific forward primer located upstream of hypothesized TE insertion position within the gene and a gene specific reverse primer downstream of the TE insertion position. genomic PCR2 (GPCR2) was performed using the same gene specific forward primer used in the GPCR1 and a TE specific reverse primer, whereas GPCR3 was performed using the gene specific reverse primer used in the GPCR1 and a TE specific forward primer. All primer sequences are listed in **Table S2**.

### RNA extraction, cDNA synthesis, RT-PCR, quantitative RT-PCR (RT-qPCR), and droplet digital PCR (dd-PCR)

Total RNA was extracted from 7 day old 30-50 female fly heads using NEB Monarch Total RNA Miniprep Kit and quantified using Thermo Fisher Scientific NanoDrop One Microvolume UV-Vis Spectrophotometer. First Strand cDNA synthesis was performed according to the NEB specified protocol using random primers, ProtoScript II (NEB), and 1 μg of total RNA input.

Quantitative PCR (qPCR) was performed using the NEB Luna Sybr-Green mastermix with different primer sequences described in Table S2. For different gene and primer combination, qPCR reaction was optimized by making a serial dilution of the original cDNA reaction ranging from 2X to 10X dilution. Relative changes in gene expression were calculated using the 2^ΔΔCt method with *Rp49* as a housekeeping gene for normalization. Briefly, the ΔCt value difference between target gene (*Dscam2* and *Bx*) and housekeeping gene (*Rp49*) was calculated for *w1118* (experimental group-TE insertion) and *OreR-MOD* (control-no TE insertion); and the difference between these two ΔCt values (dΔCt experiment- dΔCt control) was further calculated to obtain the ΔΔCt value. Relative fold change values (from experiment to control) were calculated from the exponent of 2 to the power of negative ΔΔCt value.

Droplet digital PCR (ddPCR) was conducted according to the protocol described in Yang et al [16]. Briefly, ddPCR was performed on a Bio-Rad QX200 instrument with the ddPCR Evagreen Supermix (Biorad). Copy number measurements for specific genes (Table S1) were normalized to *Rp49* using 2 ng of cDNA as input per 20 μL ddPCR for droplet generation. For genes with very high copy numbers (for example *Rp49*) that saturate the droplets, input cDNA was diluted further into the ddPCR mix prior to droplet generation. At least 12,000-15,000 droplets were generated to achieve good statistical estimation of the concentration calculated by Poisson distribution using Quantasoft Analysis Pro (Biorad).

## ACKNOWLEDGEMENTS

We thank members of the Lau lab, Reazur Rahman and Xiaoling Zhang for comments on this manuscript.

## AUTHOR CONTRIBUTIONS

M.F.A. performed the wet-lab experiments while T.T. carried out the TIDAL and TEchim bioinformatics analysis. M.F.A and N.C.L. wrote the paper with input from T.T. N.C.L. provided the direction and funding for this study.

