## Supplemental Figure S1 for "Transposable Element (TE) insertion predictions from RNAseq inputs and TE impact on RNA splicing and gene expression in *Drosophila* brain transcriptomes"

Azad, Tong et al.
Figure S1.

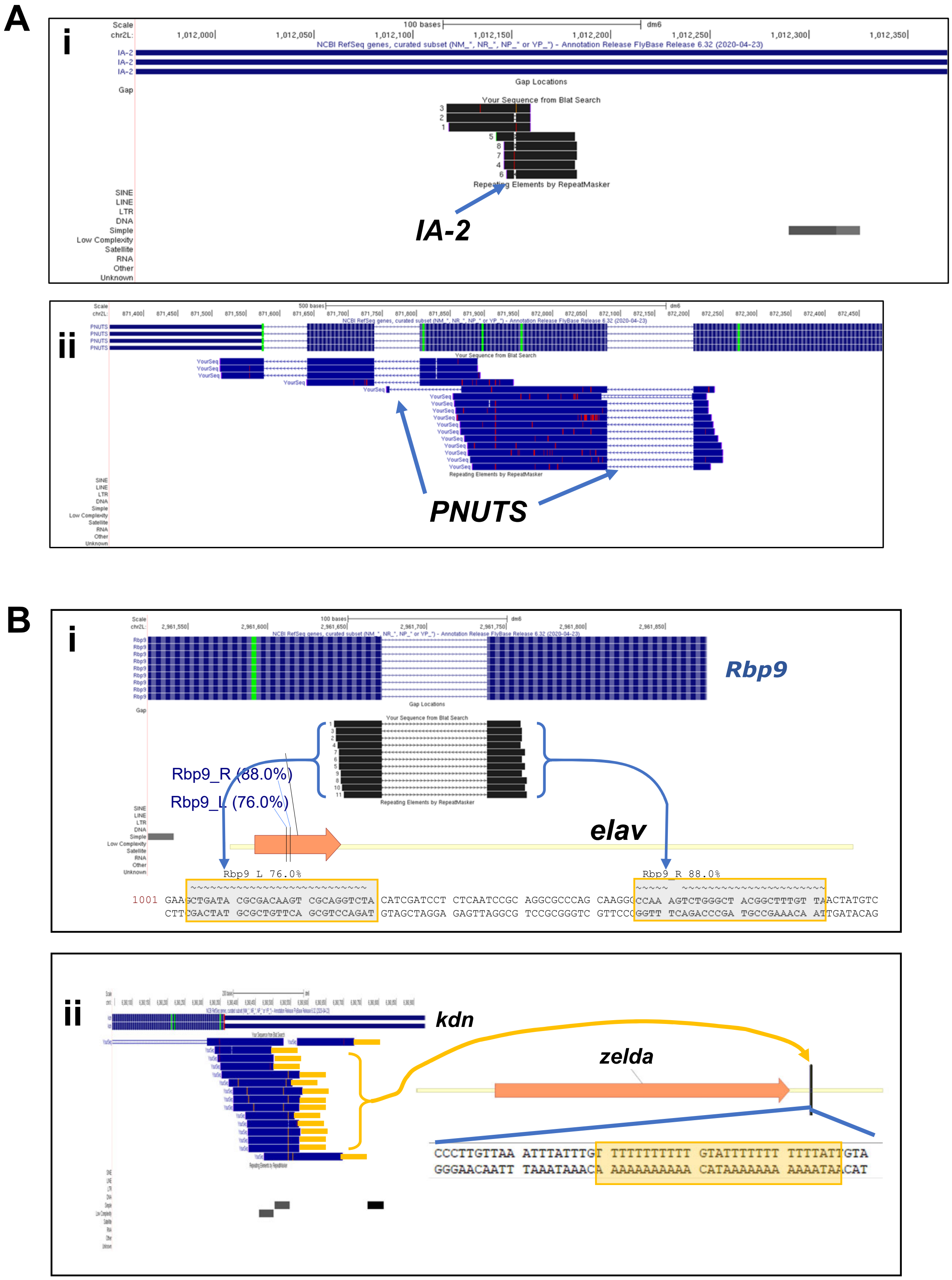


**Figure S1. Examples of why TIDAL is calling events like gene InDels/Splicing events and false two-mRNA fusion events from using RNAseq reads as inputs.**
(A) UCSC Genome browser coverage plots of RNAseq reads from the *w1118 Drosophila* midbrain of (i) a small InDel in the *IA-2* gene and (ii) intron-spanning reads of the *PNUTS* gene, which are genes loaded into TIDAL as “Immobile Genetic Elements” (IGEs). TIDAL seemed to flag these reads as a false SV call. (B) Browser plots and gene sequence snapshots demonstrate that when short split ends of longer RNAseq reads were mapped by TIDAL, sequences are commonly shared between *Rbp9* and *elav* (i) and *kdn* with another simple repeat of Poly-T’s in *Zelda* appear to cause false positive gene “fusions” being called by TIDAL.
