## Supplemental Figure S2 for "Transposable Element (TE) insertion predictions from RNAseq inputs and TE impact on RNA splicing and gene expression in *Drosophila* brain transcriptomes"

**Azad, Tong et al.**
**Figure S2.**

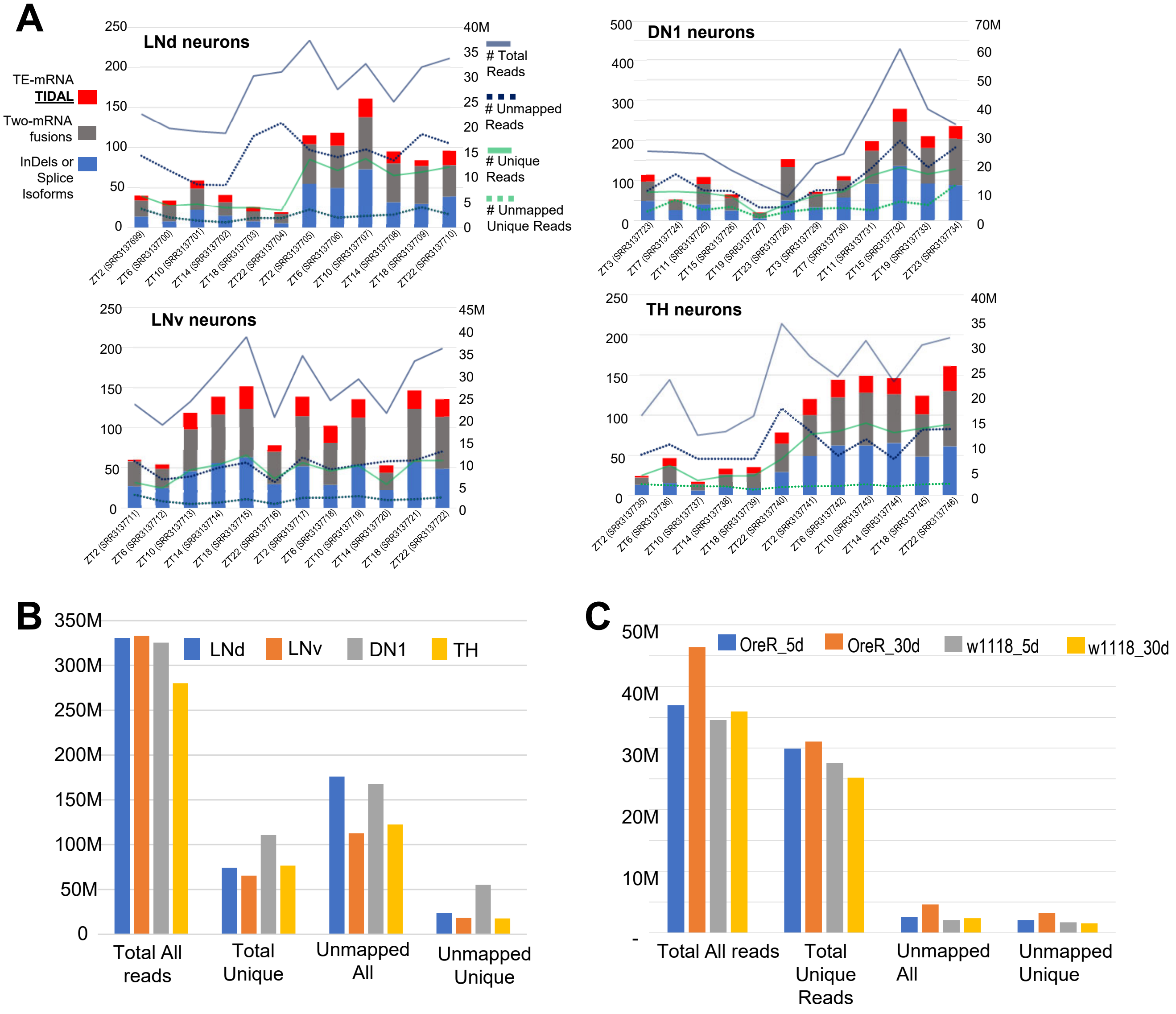


**Figure S2. Supporting analyses of *Drosophila* head/brain RNAseq and DNAseq inputs into TIDAL from results in Figures 2 and 3.** (A) Stacked bar and line graphs tallying the number of TE-mRNA fusion transcripts across the different ZT samples (library identifier in parentheses) in the different *Drosophila* neuronal types. TIDAL also reports deletion segments within the 100-control mRNA-coding genes and fusions between another mRNA with the control mRNA-coding gene. Note the two sets of legends corresponding to the different left and right Y-scale axis. The right Y-scale shows the number of total and uniquely-mapping reads as well as number of unmapped reads. (B) Read mapping statistics of the total RNA-seq data for each *Drosophila* neuron type were all Zeitgeiber timed samples were merged and then subjected to TIDAL analysis. (C) Reap mapping statistics of *OreR* and *w1118* wild-type whole genome DNA sequencing for comparison to RNAseq depths.
