## Supplemental Figure S3 for "Transposable Element (TE) insertion predictions from RNAseq inputs and TE impact on RNA splicing and gene expression in *Drosophila* brain transcriptomes"

Azad, Tong et al.
Figure S3.

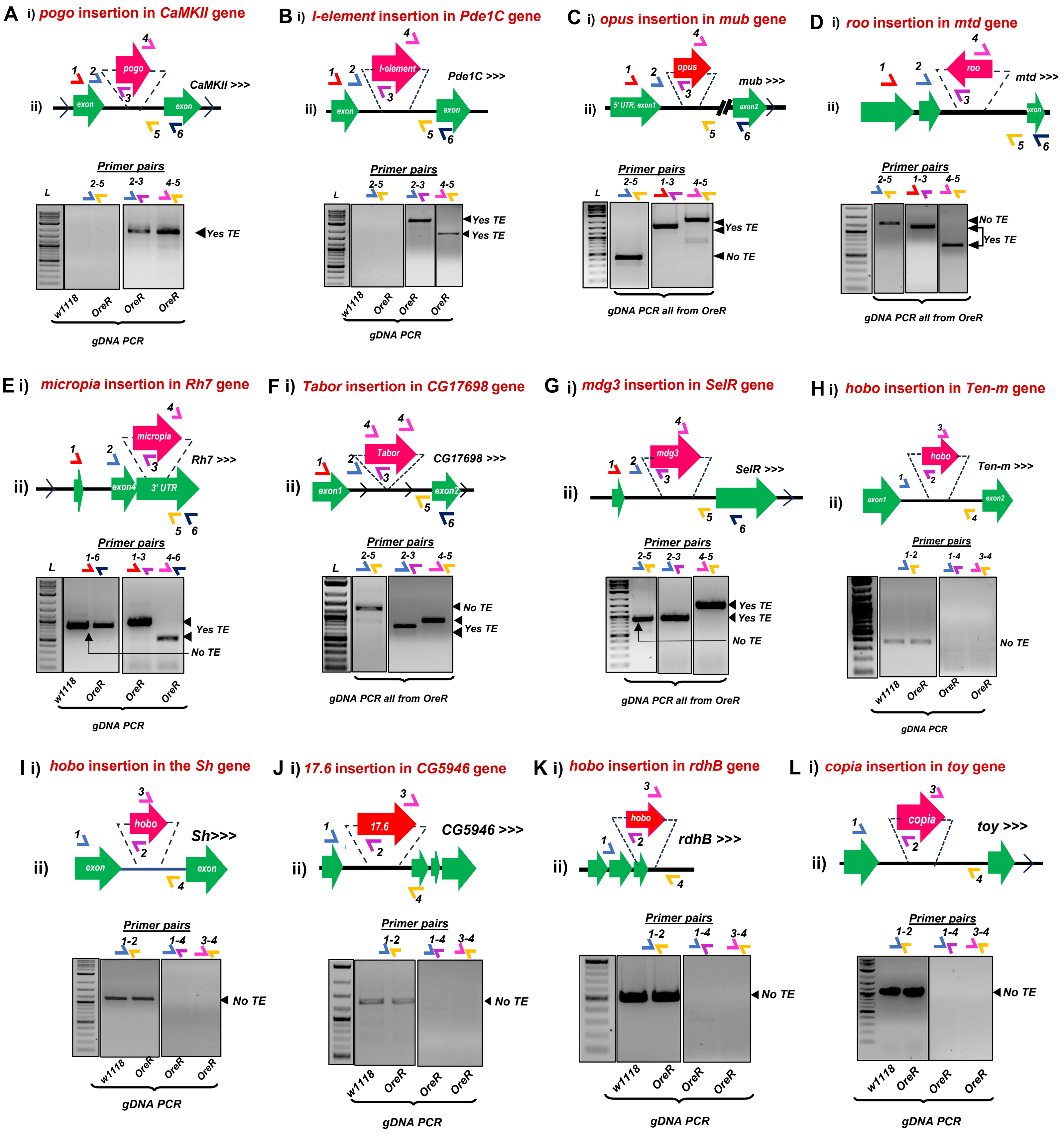


**Figure S3. Rigorous gDNA-PCR tests that confirm no TE insertion in the *Drosophila OreR* strain genome.**
Each figure panel is divided in parts (i) that is a diagram of the presumptive TE-gene splicing event proposed by Treiber and Waddell 2020, (ii) gel images of gDNA-PCR amplicons from the various sets of primer pairs illustrated in the diagram above the gel images. Solid lines around gels marked cropped gel images; dashed lines are demarcating lane sections on a single gel.

Azad, Tong et al.
Figure S3.

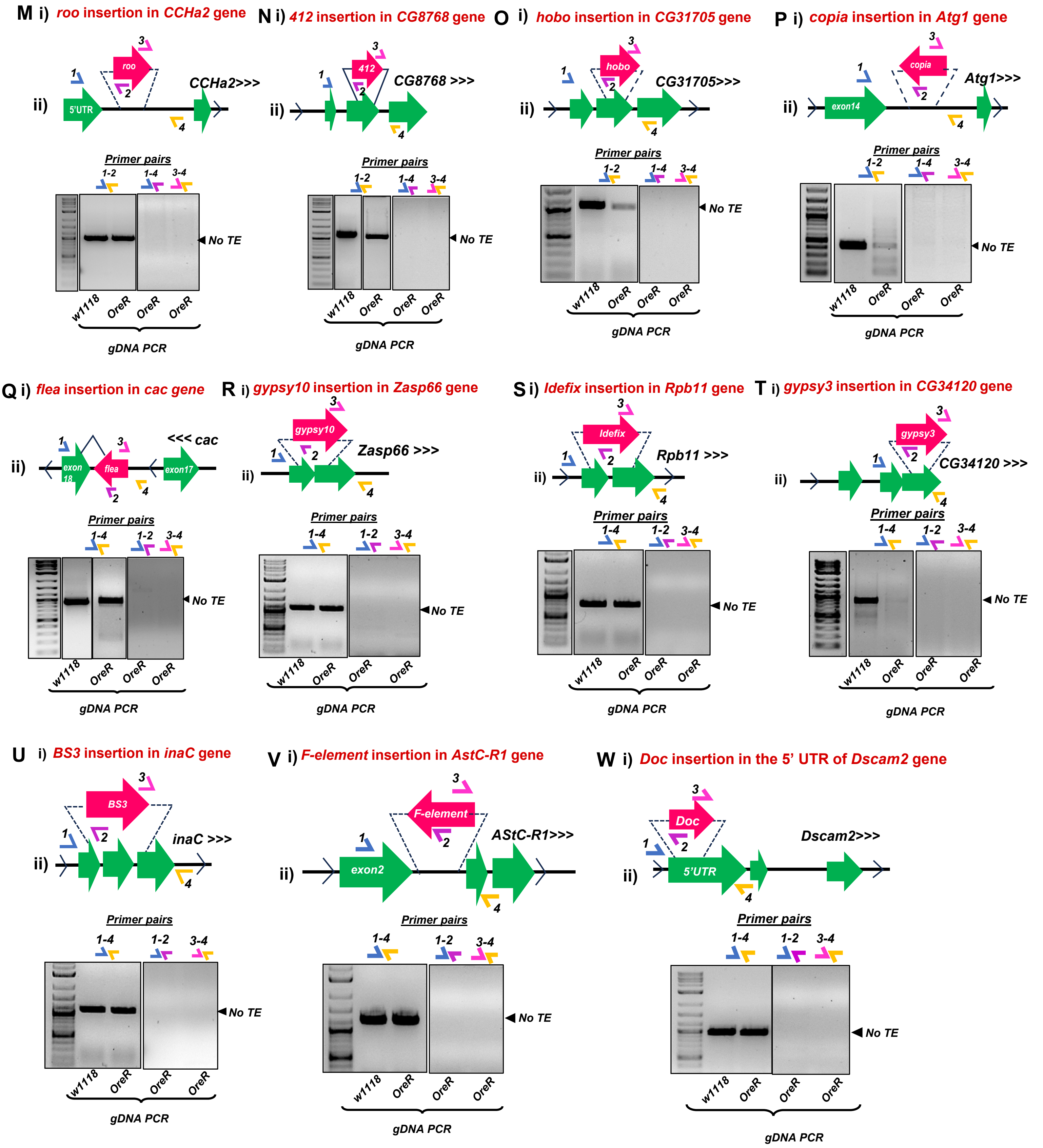


**Figure S3. Rigorous gDNA-PCR tests that confirm no TE insertion in the *Drosophila OreR* strain genome.** Each figure panel is divided in parts (i) that is a diagram of the presumptive TE-gene splicing event proposed by Treiber and Waddell 2020, (ii) gel images of gDNA-PCR amplicons from the various sets of primer pairs illustrated in the diagram above the gel images. Solid lines around gels marked cropped gel images; dashed lines are demarcating lane sections on a single gel.
