## Supplemental Figure S4 for "Transposable Element (TE) insertion predictions from RNAseq inputs and TE impact on RNA splicing and gene expression in *Drosophila* brain transcriptomes"

Azad, Tong et al.
Figure S4.

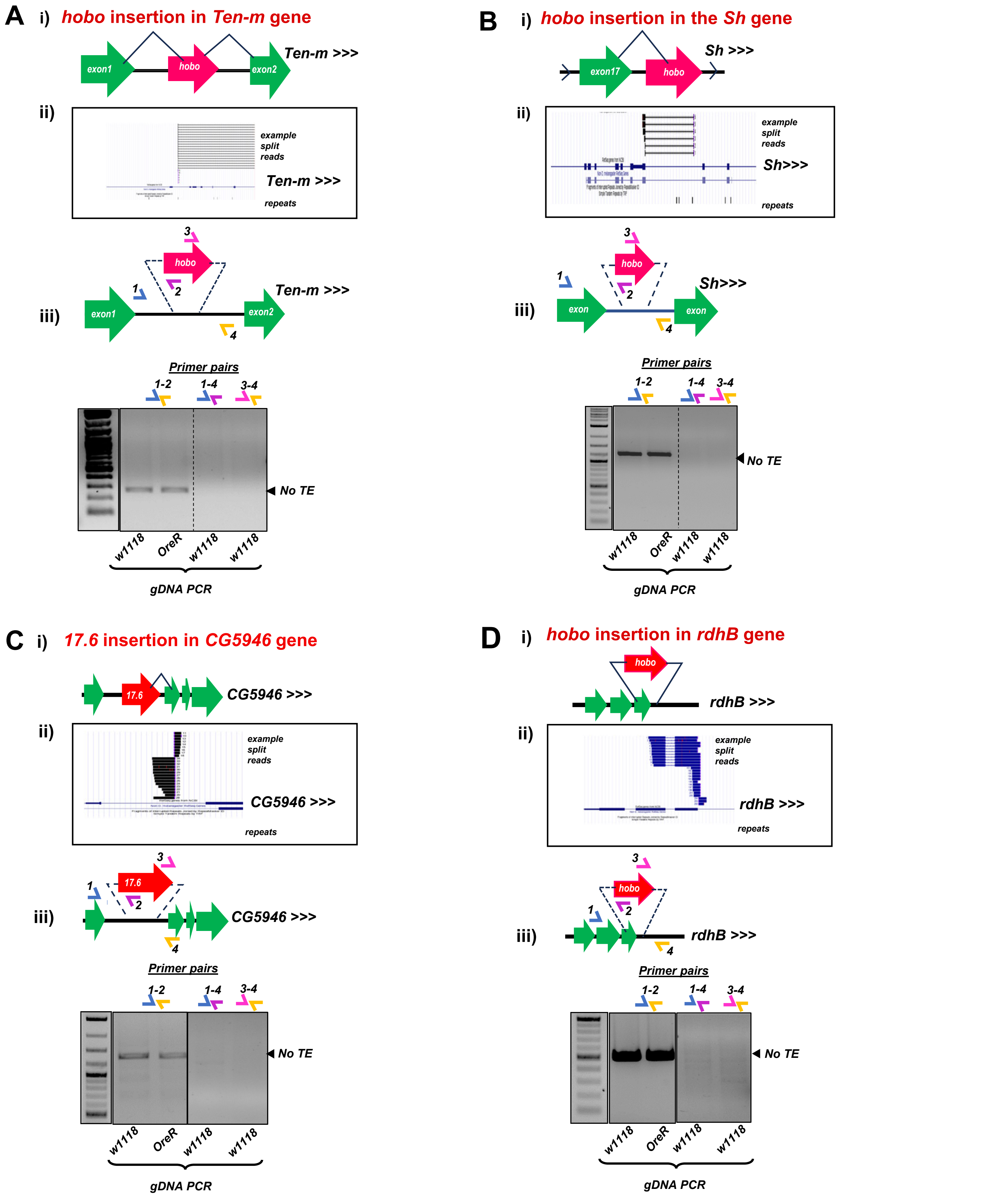


**Figure S4. Rigorous gDNA-PCR tests that cannot validate predicted TE insertions in the *Drosophila w1118* strain genome.** Panels correspond to the TE-gene pairs (A) *hobo-Ten-m,* (B) *hobo-Sh,* (C) *17.6-CG5946,* (D) *hobo-rdhB.*, (E) *copia-toy,* (F) *roo-CCHa2,* (G) *412-CG8768,* (H) *hobo-CG31705,* (I) *copia-Atg1, and* (J) *flea-cac.* Each figure panel is divided in parts (i) that is a diagram of the presumptive TE-gene splicing event proposed by Treiber and Waddell 2020, (ii) UCSC Genome Browser snapshots of the example split reads support for the TE insertion from TIDAL analysis of the *w1118* midbrain RNAseq data, (iii) gel images of gDNA-PCR amplicons from the various sets of primer pairs illustrated in the diagram above the gel images. Solid lines around gels marked cropped gel images; dashed lines are demarcating lane sections on a single gel.

Azad, Tong et al.
Figure S4

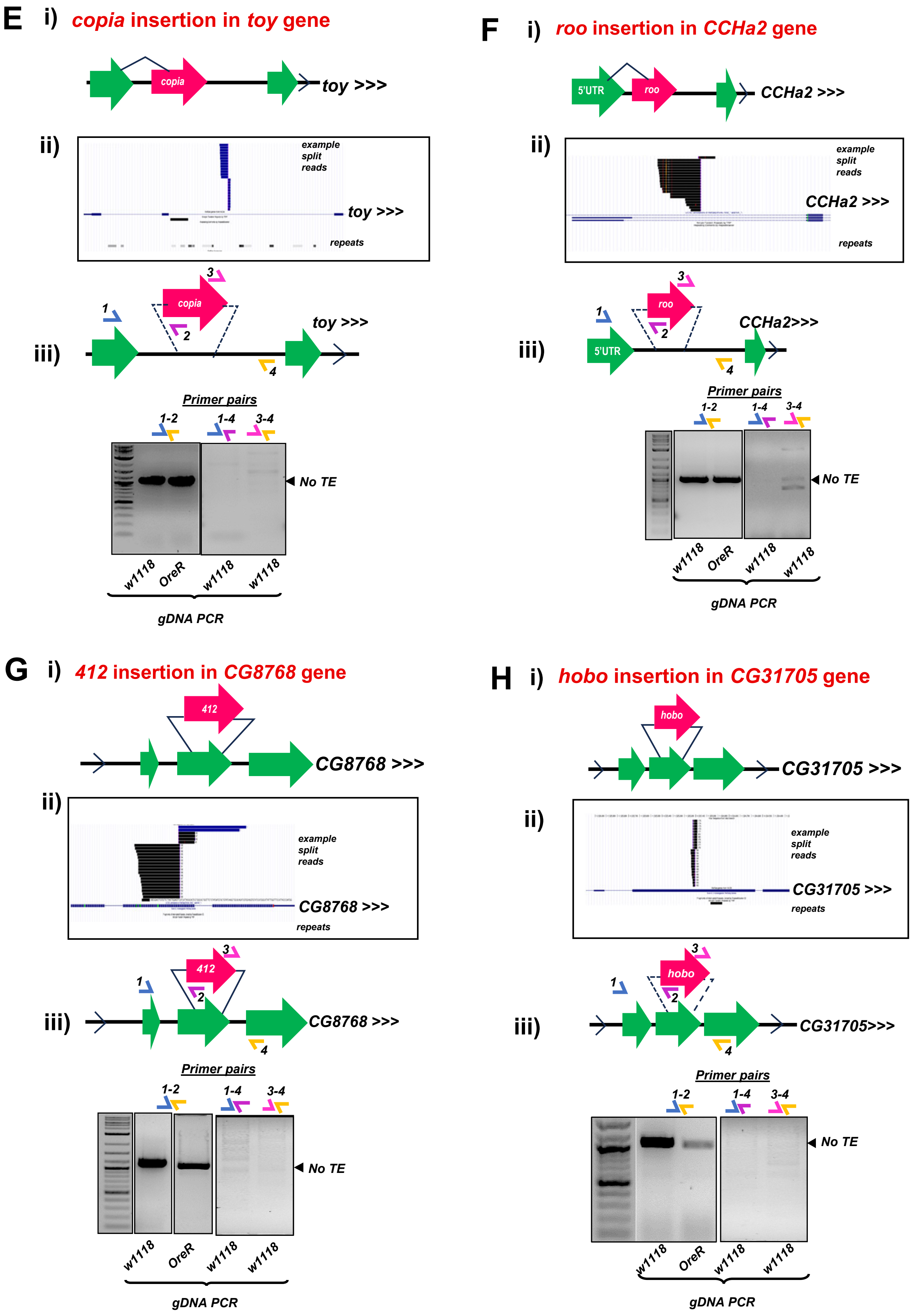


**Figure S4. Rigorous gDNA-PCR tests that cannot validate predicted TE insertions in the *Drosophila w1118* strain genome.** Panels correspond to the TE-gene (E) *copia-toy,* (F) *roo-CCHa2,* (G) *412-CG8768, and* (H) *hobo-CG31705.* Each figure panel is divided in parts (i) that is a diagram of the presumptive TE-gene splicing event proposed by Treiber and Waddell 2020, (ii) UCSC Genome Browser snapshots of the example split reads support for the TE insertion from TIDAL analysis of the *w1118* midbrain RNAseq data, (iii) gel images of gDNA-PCR amplicons from the various sets of primer pairs illustrated in the diagram above the gel images.

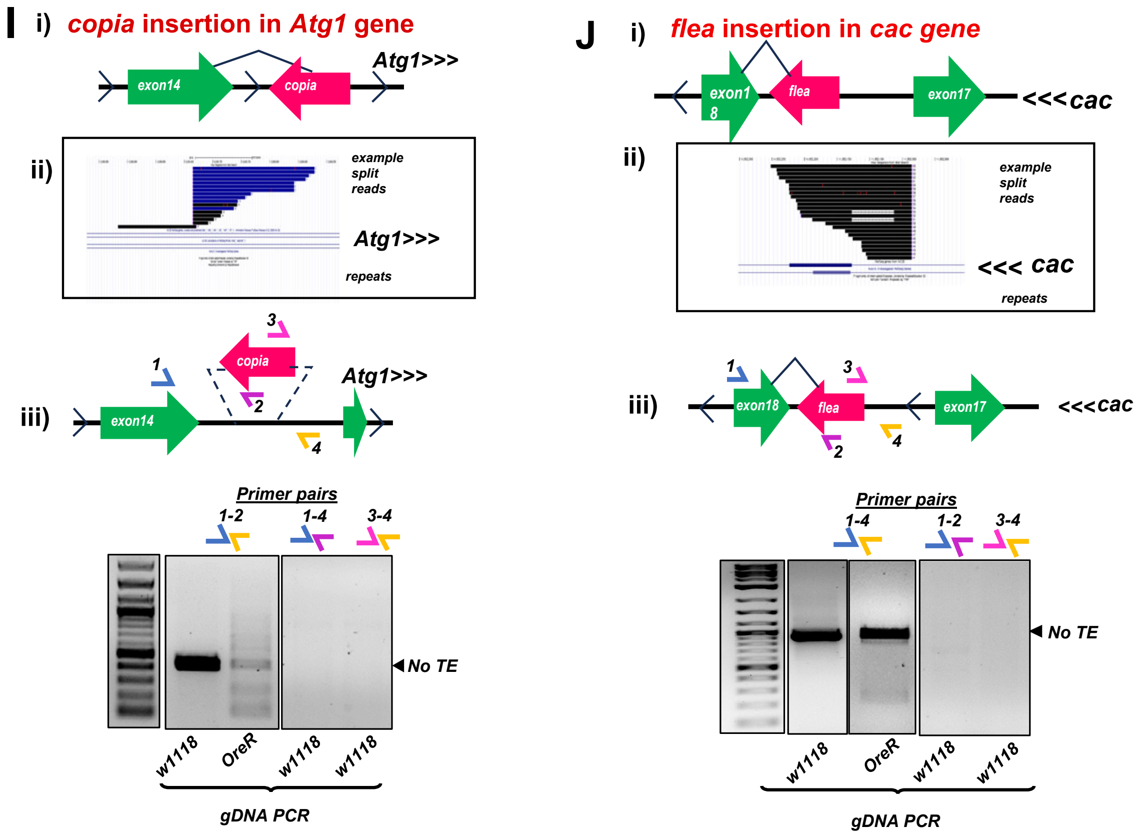


**Figure S4. Rigorous gDNA-PCR tests that cannot validate predicted TE insertions in the *Drosophila w1118* strain genome.** Panels correspond to the TE-gene pairs (I) *copia-Atg1 and* (J) *flea-cac.* Each figure panel is divided in parts (i) that is a diagram of the presumptive TE-gene splicing event proposed by Treiber and Waddell 2020, (ii) UCSC Genome Browser snapshots of the example split reads support for the TE insertion from TIDAL analysis of the *w1118* midbrain RNAseq data, (iii) gel images of gDNA-PCR amplicons from the various sets of primer pairs illustrated in the diagram above the gel images.
