## Supplemental Figure S5 for "Transposable Element (TE) insertion predictions from RNAseq inputs and TE impact on RNA splicing and gene expression in *Drosophila* brain transcriptomes"

Azad, Tong et al.
Figure S5.

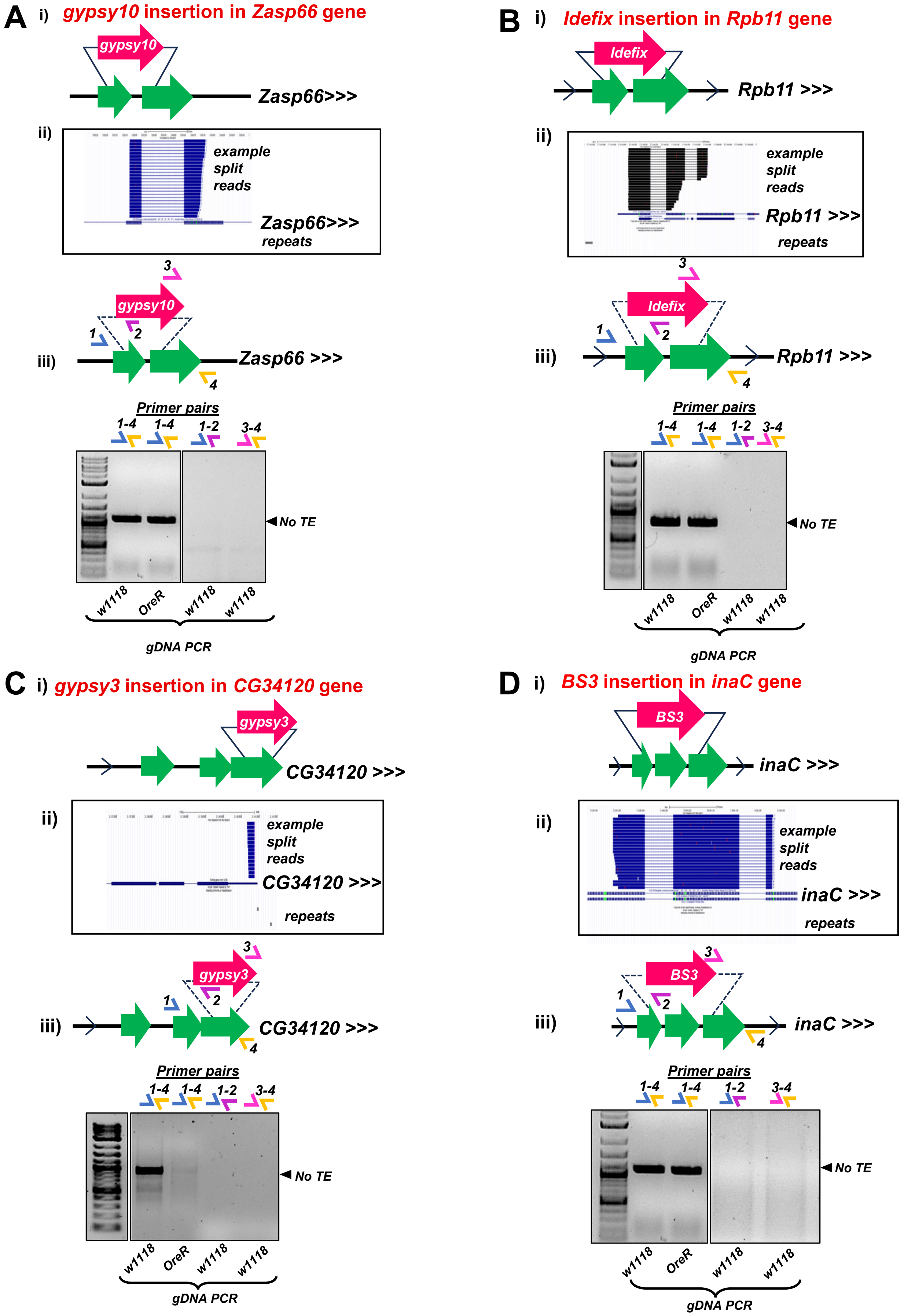


**Figure S5. Rigorous gDNA-PCR tests that cannot validate TE insertions predicted only by TIDAL in the *Drosophila w1118* strain genome.**

Panels correspond to the TE-gene pairs only predicted by TIDAL for (A) *gypsy10-Zasp66*, (B) *Idefix-Rpb11*, (C) *gypsy3-CG34120*, and (D) *BS3-inaC* Each figure panel is divided in parts (i) that is a diagram of the presumptive TE-gene splicing event (ii) UCSC Genome Browser snapshots of the example split reads support for the TE insertion from TIDAL analysis of the *w1118* midbrain RNAseq data, (iii) gel images of gDNA-PCR amplicons from the various sets of primer pairs illustrated in the diagram above the gel images.

Azad, Tong et al.
Figure. S5

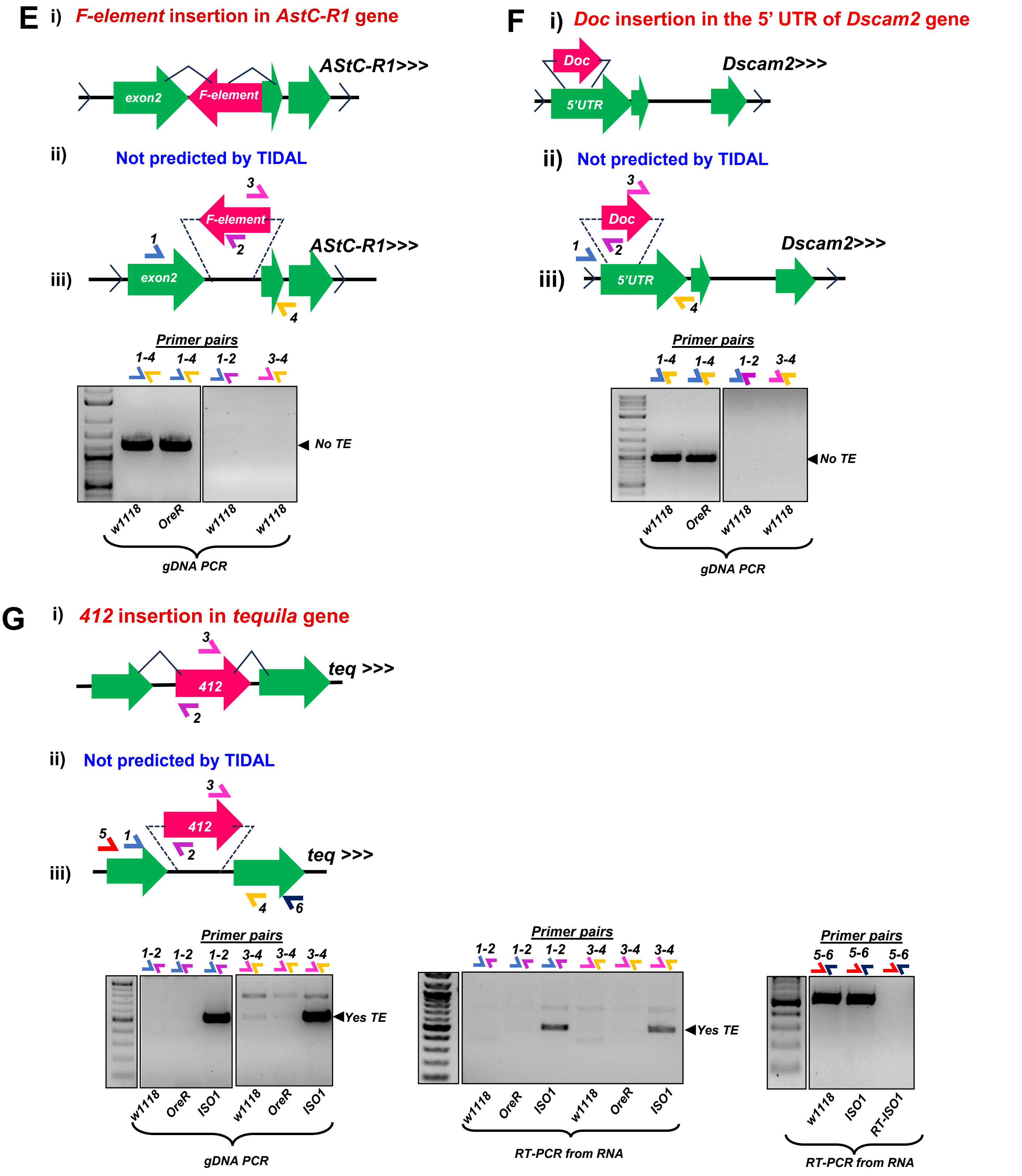


**Figure S5. Rigorous gDNA-PCR tests that cannot validate TE insertions only predicted by TEchim n the *Drosophila w1118* strain genome.**

Panels correspond to the TE-gene pairs only predicted by TEchim for (E) *F-element-AstC-R1*, (F) *Doc-Dscam2*, *and* (G) *412-Tequila.* Each figure panel is divided in parts (i) that is a diagram of the presumptive TE-gene splicing event proposed by Treiber and Waddell 2020, (ii) UCSC Genome Browser snapshots of the example split reads support for the TE insertion from TIDAL analysis of the *w1118* midbrain RNAseq data, (iii) gel images of gDNA and RT-PCR (only G) amplicons from the various sets of primer pairs illustrated in the diagram above the gel images.
