## Supplemental Table S1 for "Transposable Element (TE) insertion predictions from RNAseq inputs and TE impact on RNA splicing and gene expression in *Drosophila* brain transcriptomes"

**TABLE S1. List of Gene TE pairs evaluated by PCR in this study with detailed genomic coordinates**

| | | ***TE-Gene Pair*** | ***gDNA PCR support for TE Insertion in w1118*** | ***gDNA PCR support for TE Insertion in OreR-MOD*** | ***TE insertion position from TIDAL*** | | | ***gPCR primer set 1 tested?*** (Gene-F + Gene-R) | ***gPCR primer set 2 tested?*** (Gene-F + TE-R) | ***gPCR primer set 3 tested?*** (Gene-R + TE-F) | ***Forward primer position*** | | ***Reverese primer position*** | | ***Expected size*** | ***Observed size in w1118*** |
|---|---|---|---|---|---|---|---|---|---|---|---|---|---|---|---|---|
| Gene-TE pairs predicted in a w1118 strain derivative by Treiber & | 1 | *blood-Dscam2* | YES | NO | chr3L | 7183892 | 7184420 | √ | √ | √ | 7184411 | 7184431 | 7183660 | 718368 | 750bp | No band |
| | 2 | *opus-Bx* | YES | NO | chrX | 18550430 | 18550670 | √ | √ | √ | 18549894 | 18549914 | 3551094 | 1855111 | 1221bp | No band |
| | 3 | *I-element-Pde1C* | YES | YES | chr2L | 11860287 | 11860520 | √ | √ | √ | 11860203 | 11860222 | 1860793 | 1186081 | 613bp | No band |
| | 4 | *pogo-CaMKII* | YES | YES | chr4 | 1038286 | 1038654 | √ | √ | √ | 1037963 | 1037983 | 1038768 | 1038787 | 824bp | No band |
| | 5 | *copia-toy* | NO | NO | chr4 | 993551 | 993832 | √ | √ | √ | 993165 | 993184 | 994255 | 994274 | 1100bp | 1100bp |
| | 6 | *roo-CCHa2* | NO | NO | chr3R | 13392387 | 13392665 | √ | √ | √ | 13392841 | 13392860 | 3391900 | 1339192 | 950bp | 950bp |
| | 7 | *hobo-CG31705* | NO | NO | chr2L | 11525040 | 11525499 | √ | √ | √ | 11525733 | 11525754 | 1524726 | 1152474 | 1028bp | 1028bp |
| | 8 | *flea-cacophony* | NO | NO | chrX | 11951970 | 11952244 | √ | √ | √ | 11952372 | 11952391 | 1950263 | 1195028 | 2183bp | 2183bp |
| | 9 | *17.6-CG5946* | NO | NO | chr3L | 11819104 | 11819395 | √ | √ | √ | 11818826 | 11818847 | 1819706 | 1181972 | 898bp | 898bp |
| | 10 | *hobo-rdhB* | NO | NO | chr3R | 22352597 | 22352904 | √ | √ | √ | 22352446 | 22352464 | 2353113 | 2235313 | 689bp | 689bp |
| | 11 | hobo-Ten-m | NO | NO | chr3L | 22356882 | 22357046 | √ | √ | √ | 22356884 | 22356904 | 2357114 | 2235713 | 250bp | 250bp |
| | 12 | *hobo-Sh* | NO | NO | chrX | 17949931 | 17950074 | √ | √ | √ | 17947486 | 17947508 | 7946730 | 1794674 | 778bp | 778bp |
| | 13 | 412-CG8768 | NO | NO | chr2R | 12691622 | 12692103 | √ | √ | √ | 12692579 | 12692598 | 2691455 | 1269147 | 1143bp | 1143bp |
| | 14 | *Tabor-CG17698* | NO | NO | chr3L | 23742885 | 23743145 | √ | √ | √ | 23740772 | 23740792 | 3743295 | 2374331 | 700bp | 700bp |
| | 15 | *opus-mub* | NO | NO | chr3L | 21845047 | 21845208 | √ | √ | √ | 21844858 | 21844877 | 1845300 | 2184531 | 461bp | 461bp |
| | 16 | *roo-mtd* | NO | NO | chr3R | 5284197 | 5284467 | √ | √ | √ | 5285131 | 5285153 | 5283896 | 5283916 | 1257bp | 1257bp |
| | 17 | *micropia-Rh7* | NO | NO | chr3L | 12165142 | 12165476 | √ | √ | √ | 12165546 | 12165568 | 2164902 | 1216492 | 666bp | 666bp |
| | 18 | copia-Atg1 | NO | NO | chr3L | 12800567 | 12800901 | √ | √ | √ | 12800410 | 12800430 | 2801004 | 1280102 | 615bp | 615bp |
| | 19 | mdg3-SelR | NO | NO | chr3R | 10865047 | 10865455 | √ | √ | √ | 10865954 | 10865973 | 0865061 | 1086508 | 912bp | 912bp |
| | 20 | *Doc-Dscam2* | NO | NO | Not predicted by TIDAL | | | √ | √ | √ | 89904 | 7189924(5'UT | 3926 | 7188947(int | 924bp | 924bp |
| | 21 | *F-element-AstC-R1* | NO | NO | Not predicted by TIDAL | | | √ | √ | √ | 11077596 | 11077616 | 1076383 | 1107640 | 1233bp | 1233bp |
| | 22 | *412-Tequila* | NO | NO | Not predicted by TIDAL | | | √ | √ | √ | 9076674 | 9076694 | 9084647 | 9084666 | 7992 | No band |
| Gene-TE pairs predicted in the w1118 | 23 | *gypsy3-CG34120* | NO | NO | chrX | 21140649 | 21140805 | √ | √ | √ | 21139880 | 21139899 | 1140880 | 2114090 | 1021bp | 1021bp |
| | 24 | *gypsy10-Zasp66* | NO | NO | chr3L | 8634596 | 8634742 | √ | √ | √ | 8634080 | 8634100 | 3635117 | 8635140 | 1060bp | 1060bp |
| | 25 | *BS3-inaC* | NO | NO | chr2R | 16901112 | 16901335 | √ | √ | √ | 16900275 | 16900294 | 6901335 | 1690135 | 1080bp | 1080bp |
| | 26 | *Idefix-Rpb11* | NO | NO | chr2L | 17409980 | 17410152 | √ | √ | √ | 17410447 | 17410466 | 7409609 | 1740962 | 838bp | 838bp |
