## Supplemental Table S2 for "Transposable Element (TE) insertion predictions from RNAseq inputs and TE impact on RNA splicing and gene expression in *Drosophila* brain transcriptomes"

### Table S2. List of oligonucleotides used in this study

| genomic PCR Primer list | |
|---|---|
| Dm_Dscam2_Doc-Fw | CTGTTCGAGGCTCGAAGGATA |
| Dm_Dscam2_Doc-Rv | ATTCTTGGCCGACTCTTTTCTC |
| Dm_Doc-Internal-Fw | CACATTGTCGCTGAGAACGTAT |
| Dm_Doc-Internal-Rv | CATATCTGGGAGTTCATCTGGA |
| Dm_AstC-Fw | ACAGCAATCTGCGTCAGCAAG |
| Dm_AstC-Rv | TACCACACCATATGCAAAGCG |
| Dm_CCHa2-Fw | AGCCCAGTTCGGCTAATCAG |
| Dm_CCHa2-Rv | GTGCAGATAACGACCAGTAGCA |
| Dm_roo-LTR-Fw | AGGTGACATGAGAATCGCATC |
| Dm_roo-LTR-Rv | GTGCACACTACATGAGTCAGTC |
| Dm_toy-Fw | AGGCAAAAACGCAGCGTAAG |
| Dm_toy-Rv | GTTGGTGTAAGAAGGGGCGT |
| Dm-Copia-Fw | GTGAGTAGGTCGTGGTGCTG |
| Dm-Copia-Rv | ACCAGCACCACGACCTACTC |
| Dm_Bx-Fw | TGAGGACGATACAGCACACAC |
| Dm_Bx-Rv | ACTTGCATCAGGGTTCTCGG |
| Dm_opus-Fw | CCATCGCTCTTACGGTCCAA |
| Dm_opus-Rv | ACGGCACTTGTTCCGAATGC |
| Dm_Rpb11-Fw | AGCTAGGCGTTCTAGCTACT |
| Dm_Rpb11-Rv | TATTGAATGCGATTCGTGGC |
| Dm_CG34120-Fw | TCTCCGAACTCTTCGCCATC |
| Dm_CG34120-Rv | CTTCAGACAGATCCCATTAGGA |
| Dm_inaC-Fw | ATGAGCTGTATGCCGTGAAG |
| Dm_inaC-Rv | TCCGTTGGAGTTAAGTCCGTC |
| Dm_gypsy3-Fw | ACTAAGTTAACCGGACTGATCGTC |
| Dm_gypsy3-Rv | GAGCCATCGCTCATTAACCAACA |
| Dm_Zasp66-Fw | ACGAGCAACAGCTGATCAAGC |
| Dm_Zasp66-Rv | CATTGTATCGAGCTAAGCATAGGA |
| Dm_CG5946-Fw | CTGCTGAATAAGAAGTCCACGA |
| Dm_CG5946-Rv | TAGCCATGACGTCACAGCA |
| Dm_17-6-Fw | AGCGGCACTTAGCCATTCTT |
| Dm_17-6-Rv | AGGAGAAGCCTCTGTGCTTG |
| Dm_CG17698-Fw | TCAATCGATATGCTAACAAGAACGC |
| Dm_CG17698-Rv | ACGGACATGTACAGATCGACTC |
| Dm_Tabor-LTR-Fw | ATTCAGACCAGAAGTGCAGAGTC |
| Dm_Tabor-LTR-Rv | GAAGGCTCTTTGACGACTCCT |
| Dm_CaMKII-Fw | CTGACATGATCGACTCAGCTA |
| Dm_CaMKII-Rv | TCACTAGCCGAGTCGACTTG |

| | |
|---|---|
| Dm_pogo-Fw | TGCATCGATAGTTAGCTGCATC |
| Dm_pogo-Rv | GCTTAGCTGCCTCGAGTACT |
| Dm_rdhB-Fw | AGCTCGTCGAGTGGAACTC |
| Dm_rdhB-Rv | GAACACTTTCTACGGAATTGGCT |
| Dm_Ten-m-Fw | AGTTCTGTAAGCGCAGATAAGAC |
| Dm_Ten-m-Rv | CTGTCCTTAAAGGATTTCGCAG |
| Dm_Sh-Fw | ACTGCTACTTATGAACCGTAACG |
| Dm_Sh-Rv | TGAATTGCGATCCGAATGCG |
| Dm_hoboFw | ATCGTTGACTGTGCGTCCACT |
| Dm_hobo-Rv | TGCGCACCCGAATCAATACG |
| Dm_Pde1c-Fw | CCGAAGCCATATGGCAATGA |
| Dm_Pde1c-Rv | GTCTAACTATAGCTGTACAGACCA |
| Dm_I-element-Fw | ACGCTGGATAGGAGTTGAGATG |
| Dm_I-element-Rv | TGAGAGGCACGACTTATCTCTTC |
| Dm_Rh7-Fw | AGACCGAACTGAAATCAGCAATG |
| Dm_Rh7-Rv | ACCACCGGATTAGACATCGAG |
| Dm_micropia-Fw | ATCGTGGAGAAGCCAAAGCA |
| Dm_micropia-Rv | GTCCGTCCGTCCATATCAGC |
| Dm_Atg1-Fw | TAGGTGCAATGGCCTCTCAAC |
| Dm_Atg1-Rv | ACAGCGTAAATACACGGAAGTC |
| Dm_mub-Fw | ATCAGCCGACCGAGTTTGAC |
| Dm_mub-Rv | ATGGCAACAAGAACTGCTCG |
| Dm_opus-Fw | CACTCAGGGTGAGGGGTCAA |
| Dm_opus-Rv | TCGAGACTGGGACCTCTTCT |
| Dm_CG8768-Fw | AGGCATGCTCTGATAGGTAC |
| Dm_CG8768-Rv | TGCGCTTGGTACTAACATCTA |
| Dm_SelR-Fw | TGTTCCACTGCCCAGAGTTC |
| Dm_SelR-Rv | CCAGGAACTGTAGACTCGCA |
| Dm_mdg3-Fw | TAATCGACTCGCCACTCTGC |
| Dm_mdg3-Rv | TAGCCGCCGTTTACAGAAGT |
| Dm_mtd-Fw | TGCACGTATGCTCTCGTATCAAC |
| Dm_mtd-Rv | AGCATTCCAACTCGAAGTTGC |
| Dm_roo-LTR-Fw | AGGTGACATGAGAATCGCATC |
| Dm_roo-LTR-Rv | GTGCACACTACATGAGTCAGTC |
| Dm_cac-Fw | ATAGAGTTTTAGCCATGCAACTC |
| Dm_cac-Rv | TGAGCTAGGCTATTAGTAGTTTGA |
| Dm_flea-LTR-Fw | AGTGGAAGTCAGCGTTGCAG |
| Dm_flea-LTR-Rv | CGACAATCATGTTGCTGCTCA |
| Dm_CG31705-Fw | ACATATGTATGTGGCAGGCATG |
| Dm_CG31705-Rv | ACTGGATGGTAAGATGCTCCTGC |
| Dm-Teq-Rv | AGGTGGGCAGGAAATGTCAC |

| | |
|---|---|
| Dm-Teq-Fw | AGAGATCATATGCTGTCCCAC |
| Dm-412-Rv | ATTTGGTCGGCGTGTGAATG |
| Dm-412-Fw | AAGTGCATTGCCCACTCGAA |
| Dm_BloodLTR-Fw | CTAAGTCAGCATCCCCACGC |
| Dm_BloodLTR-Rv | CTGTAATAAACCAATATATGCC |
| Dm_Dscam2-Fw | ACTTTCGAATGCGACAATGGC |
| Dm_Dscam2-_Rv | CACCGAATTGCTGGCAATACAGC |
| Dm-blood-cds3-FW1 | ATGCGAATATCACCAAGCGAAC |
| Dm-blood-cds3-FW2 | CTATAGAAGGAGGCCACACTG |
| Dm-blood-cds3-FW3 | CCAGATGACCAAGACAATCTCA |
| **qRT-PCR and ddPCR primer list** | |
| Dm_Dscam2-Ex1-Fw | ATTGTACAGCTCACTGCCCAC |
| Dm_Dscam2-Ex2-Rv | GCTTCGAACGGTTCACTGCA |
| Dm_Dscam2-Ex3-Fw | AGGAATTGGTACGAGTGGTCTC |
| Dm_Dscam2-Ex4-Rv | AGATTGTGGATGAGCAGCTC |
| Dm_Dscam2-Ex4-Fw | AGAGCGATGAGTCGCAGTC |
| Dm_Dscam2-Ex5-Rv | GTATTCGGGCGAAGGACAG |
| Dm_Bx-Ex1-Fw | GCAACAGTAAGAGCCTTGTGC |
| Dm_Bx-Ex2-Rv | ACCGGGTTCGGTTTTGGTAA |
| Dm_Bx-Ex2-Fw | AGCCTTCGAGATGGTGATGC |
| Dm_Bx-Ex3-Rv | ATTGGCCATCGATGCGAAGAC |
| **RT PCR Primer List** | |
| Dm_mtd-UTR-Fw1 | TCTCCACGATTCTGCGATTCTTC |
| Dm_mtd-Exon3-Rv1 | TGTTGTGAGGTGCTGAAGATCAG |
| Dm_mub-UTR-Fw1 | AGTGCATAATCCTTCAACAGC |
| Dm_mub-exon-Rv1 | ATCCTCGTGTTTGATGGACG |
| Dm_Rh7-UTR-Fw1 | AGACCGAACTGAAATCAGCAATG |
| Dm_Rh7-UTR-Rv1 | ACCACCGGATTAGACATCGAG |
| Dm_Pde1C-UTR-Fw1 | TGGAAGCCATTGCAGCTAAC |
| Dm_Pde1C-exon-Rv1 | TAGTTGGTTGCATTCGGCGATG |
| Dm_SelR-Exon-Fw1 | ACCAGGATCTGTTCAGCTCG |
| Dm_SelR-Exon-Rv1 | GAATTGATGCAGTAGCGCTTGC |
| Dm_CaMKII-exon-Fw1 | AGGAACCGTTGATCGCAGTAC |
| Dm_CaMKII-exon-Rv1 | TCTACAAGGTTACCCAGTGCTTC |
| Dm_CG17698-FW1 | ACAAGTCCTAAGGAAAGAAATGC |
| Dm_CG17698-RV1 | ATGAGTGGAATCTTCTTCGGAG |
